# Mapping the phenotypic landscape of a transcriptional repressor using Deep Mutational Scanning and Growth-based Quantitative Sequencing

**DOI:** 10.64898/2025.12.11.693801

**Authors:** Zachary Jansen, Xuan Le, Qiyao Wei, Devon L. Kulhanek, Nina Alperovich, Olga Vasilyeva, Andrew R. Gilmour, David Ross, Ross Thyer

## Abstract

CymR is a TetR-family transcriptional repressor that recognizes a well-defined operator sequence in the promoter P*_cymRC_*. The native ligand cumate and several structurally related aromatic acids bind at an allosteric site and induce a conformational change in CymR, resulting in release from the DNA operator and de-repression of the promoter. The amino acid residues that contribute to these core functions have not been mapped, nor has the protein been subjected to extensive mutagenesis to modify its function. Here, for the first time, we integrate Deep Mutational Scanning (DMS) with Growth-based Quantitative Sequencing (GROQ-Seq) to evaluate a comprehensive phenotypic landscape of CymR variants, including single amino acid insertions and deletions. We measure this library across a concentration gradient of small molecule inducers to construct an induction curve for all library members. From this analysis, we identify amino acids throughout the protein that are essential for repressor function and discover several mutations that improve the sensitivity of CymR to the ligand perillic acid. In addition, rarely investigated insertion mutants are revealed to be a key driver of novel phenotypes, including several regions of CymR where insertions result in an inverted phenotype and the isolation of variants exhibiting an unusual band-stop phenotype.

**Graphical Abstract:** 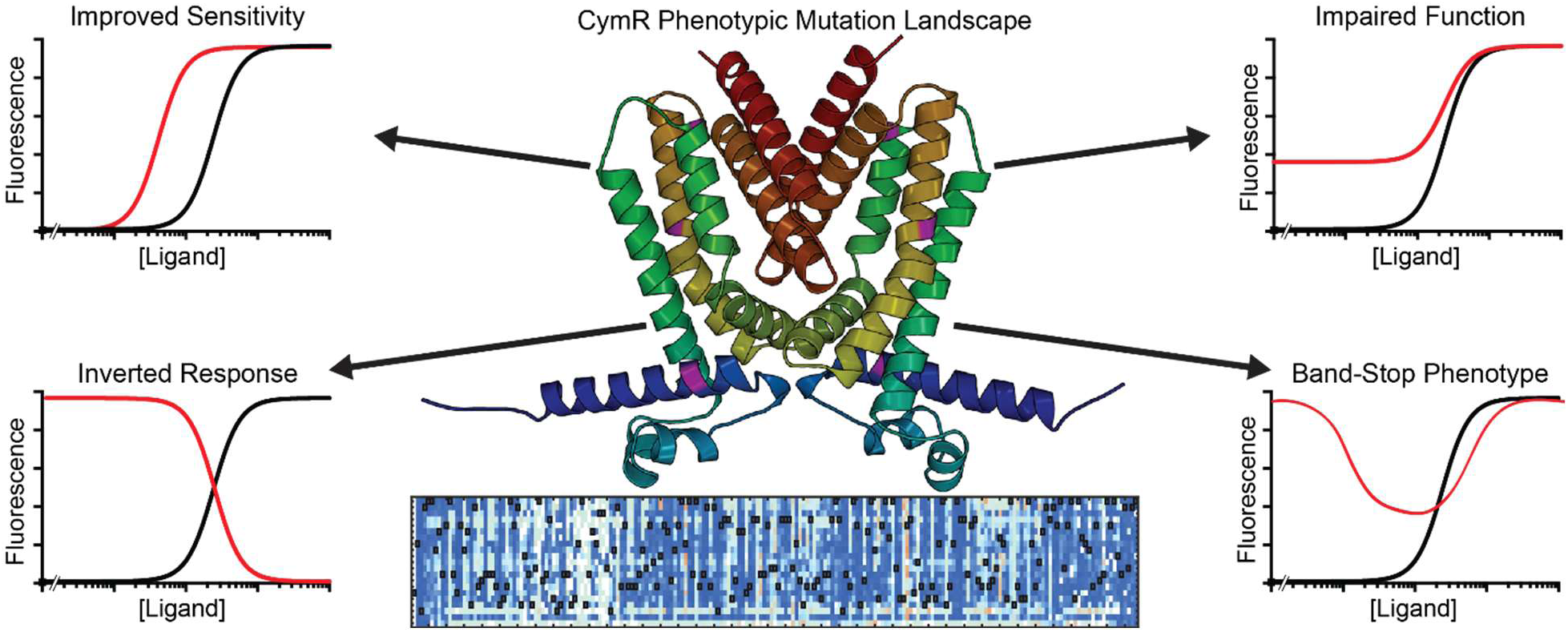

## Introduction

Transcription factors are a cornerstone of gene regulation, exerting fine control of gene expression in response to diverse chemical and physical stimuli, as well as underpinning the engineered genetic circuits that drive synthetic biology applications^1–3^. TetR-family transcriptional repressors comprise a large family of transcriptional regulators that repress transcription when bound to their cognate operator sequences. TetR-family transcriptional repressors comprise two discrete domains, a helix-turn-helix DNA-binding domain and a ligand-binding domain composed of a helix bundle. Small molecule ligands can reversibly bind within a cleft in the ligand binding domain, inducing an allosteric conformational change in the protein structure that is transduced to the DNA-binding domain such that it disfavors binding to the DNA, thereby removing the steric block on transcription^4^. As they interact solely with the DNA operator and not the host’s transcriptional machinery, TetR-family transcriptional regulators are amenable to heterologous expression, making them particularly well-suited for regulating synthetic genetic circuits^5–11^. Efforts to develop improved variants of TetR have identified several useful phenotypes, including an inverted phenotype which has been adapted as the reverse tetracycline-transactivator (rtTA) and is widely used in mammalian cells^12–14^.

CymR is a TetR-family transcriptional repressor native to *Pseudomonas putida* that binds to a defined operator sequence in the promoter PcymRC^15^. Cumate, a metabolite of the alkyl-substituted aromatic hydrocarbon p-cymene, is the native ligand. However, structurally similar aromatic acids can also bind and induce a conformational change in CymR, including di-methyl amino-benzoic acid (DMABA) and perillic acid (PA). Possibly due to the non-planar structure of PA, CymR is markedly less sensitive to this ligand.

CymR is widely used to control genetic circuits in synthetic biology, however, no empirically determined structure is available for CymR and there have been only two mutagenesis studies probing its function^16–19^. In one report, a variant of CymR was identified which has an inverted phenotype similar to the widely used reverse-TetR. In the second example, salicylic acid was discovered to have an antagonistic effect on CymR induction and a variant was identified which reduces this effect^20,21^. In both examples, variant libraries were generated using error-prone PCR, resulting in both sparse and biased sampling per residue while yielding multiple mutations per gene. Thus, despite its widespread usage, CymR has never been systematically investigated to map the functional importance of each residue in the protein, nor which amino acids or regions may contribute to novel functions.

Two key technologies are needed to comprehensively map the phenotypic landscape of a transcriptional repressor: methods to generate mutational diversity with low bias and wide sequence coverage, and measurement tools capable of capturing a complete range of sensor phenotypes^22^. Deep Mutational Scanning (DMS) is well-suited to studying global protein properties such as allostery, as a defined set of mutations can be generated with minimal bias and no prior information about the functional importance of particular residues^23^. Construction of the DNA libraries needed for comprehensive DMS has become substantially more accessible with the development of oligonucleotide pools of sufficient length and complexity to span genes within a practical number of fragments, along with computational tools to generate the required sequences for the oligonucleotide subpools^24–27^. This method allows researchers to rapidly generate protein libraries with low bias and high sequence coverage, and importantly, opens the door to sampling mutations not commonly observable using PCR-based mutagenesis or continuous evolution methods, such as insertions and deletions of whole codons^28^.

The recently described <u>gro</u>wth-based <u>q</u>uantitative <u>seq</u>uencing (GROQ-Seq) is a powerful measurement tool that can provide in-depth data on a large protein sequence space by quantifying protein function on the basis of cell fitness^29,30^. GROQ-Seq can evaluate an entire library of tens to hundreds of thousands of variants, sequencing the variants before and after selection to quantify enrichment across multiple time points^29,30^. This process can be implemented in a single well of a 96-well plate, and thus is easily scalable, enabling the library to be evaluated under numerous conditions in a single experiment. In coordination with previously evaluated calibration variants, the phenotype of each variant in each condition tested can be quantified based on changes in abundance within the library population measured by DNA sequencing. For an allosteric transcriptional regulator such as CymR, evaluating variants across a range of small-molecule inducer concentrations generates a full induction curve for each member of the library from a single experiment. This workflow shares some characteristics to the widely used Sort-Seq method, however, evaluating the population in aggregate instead of through individual cell measurements avoids challenges with cell-to-cell variability and the task of assigning bins to classify variants^31,32^.

In this work, we use GROQ-Seq to evaluate a DMS library including near complete coverage of single amino acid substitutions, insertions, and deletions for the first time to fully map the phenotypic effects of single amino acid mutations in the widely used transcriptional repressor CymR. We evaluated the library of variants across a gradient of ligand concentrations, measuring a full induction curve for every variant present in the library in parallel to reveal how each mutation impacts the function of CymR. Our data maps which amino acids are compatible with each position of CymR, revealing the residues important for repressor function, and identifies regions where insertion mutations are tolerated or deleterious. Analysis of the full phenotypic landscape enables the identification of mutations that improve the sensitivity of CymR to the non-standard ligand PA. Unexpectedly, it also reveals that mutations leading to an inverted response are common, widely distributed throughout the protein sequence, and are most often accessed via amino acid insertions. Investigation of the inverted phenotype reveals that it can be further enhanced through combinatorial mutagenesis. A novel band-stop phenotype in a transcription factor is also identified for only the second time, the first time where this phenotype is conferred by a single mutation.

## Methods

### Molecular Biology

PCR amplification was performed using a high-fidelity DNA polymerase. Products were analyzed by agarose gel electrophoresis and purified using the Zymo Clean & Concentrator kit. Backbone PCR products were digested with DpnI to eliminate template plasmid background. All plasmids described in this work were constructed using either Gibson Assembly or a previously reported hierarchical Golden Gate assembly system^33^. Gibson Assemblies were performed with a 1:3 ratio of backbone:insert DNA and incubated at 50 °C for 2 hours. Golden Gate assembly reactions were performed by combining 20 fmol of each DNA fragment, T4 DNA ligase, T4 DNA ligase buffer, and the appropriate restriction enzyme. Assemblies were transformed into *E. coli* DH10B or DB3.1 using chemical transformation. Cultures were incubated at 37 °C in Terrific Broth supplemented with antibiotics at a concentration of 33 ug/mL for chloramphenicol and 50 ug/mL for ampicillin. Annotated sequences from plasmids used in this work are included as supplementary GenBank files.

### Library Construction

A modified version of the SPINE tool was used to design the mutagenic oligonucleotide library tiles for *cymR*. Briefly, the *cymR* coding sequence was fragmented into seven library tiles. Each tile was ordered from Agilent Technologies as a pool of 150 nt synthetic ssDNA fragments encoding type IIS restriction sites at both ends and a unique pair of amplification tags. Tile-specific primers were used to amplify each library tile from the total pool to generate dsDNA fragments suitable for assembly. For each library tile, a corresponding backbone fragment encoding the remainder of the *cymR* gene along with the core elements of the plasmid was amplified by PCR from a wild-type parental plasmid. The concentration of template DNA was minimized to reduce carryover of the wild-type sequence into the library population, and PCR products were digested with DpnI to further minimize template contamination. Approximately 300 ng of backbone DNA was prepared for each assembly. Each tile fragment was assembled with its matching backbone by Golden Gate assembly. Each 20 µL reaction contained 300 ng of backbone PCR product, a 1:1.25 molar ratio of insert DNA, 1 µL BsaI, 1 µL T4 DNA ligase, 2 µL CutSmart buffer, and 1.25 mmol/L ATP. Reactions were incubated using a thermocycler at 37 °C for 5 minutes and 16 °C for 2 minutes, repeated for 64 cycles, followed by incubation at 37 °C for 30 minutes and enzyme inactivation at 85 °C for 15 minutes. The resulting assemblies were pooled and purified by ethanol precipitation to maximize DNA recovery.

The pooled library was transformed into *E. coli* DH10B using electroporation. Transformation efficiency was determined by dilution plating (10^6^ transformants) and exceeded more than 10-fold the possible number of CymR variants in our library, ensuring sufficient coverage. Library plasmid DNA was recovered from liquid cultures to serve as the template for a subsequent barcoding reaction. DNA barcodes were introduced by PCR using primers encoding 18 degenerate bases and a fixed internal homology region to facilitate recircularization by Gibson Assembly. Barcodes were introduced into a designated region downstream of the *cymR* gene flanked by terminator sequences to reduce the possibility of cryptic barcode sequences influencing other plasmid elements. The barcoded library was purified by ethanol precipitation and transformed by electroporation again exceeding 10-fold coverage (10^6^ transformants) of the theoretical library size.

### Selection of Normalization Controls and Protein Function Calibration Ladder for GROQ-Seq

Two different types of reference ‘spike-in’ variants were used to calibrate the GROQ-Seq DMS data. First, normalization controls which were used to normalize the barcode read counts in each sample to account for sample-to-sample variability in DNA extraction efficiency and/or barcode PCRs, and second, a set of 16 CymR variants, with a range of different functional responses, which were used as a calibration ladder to convert enrichment values (derived from barcode sequencing counts) to functional values (the expression level of genes regulated by the CymR transcription factor), providing a full induction curve for every variant in the library. An extended discussion of how these were selected and used is provided in the **supplementary methods,** including **Tables S1 and S2**.

#### GROQ-Seq DMS Assays

The growth-based quantitative DMS assay was performed using a protocol similar to the one used for previous work with the lac repressor (LacI) and the RamR transcriptional regulator^29,30^. An extended description of the methods used is provided in the **supplementary methods**.

#### Fluorescence Assays

All fluorescence assays were performed in *E. coli* strain DH10B grown in M9 medium supplemented with 2.5 g/L yeast extract (M9YE) and 0.5% v/v glycerol, unless otherwise specified. Transformants were selected in biological triplicates into 96-well deep-well plates and incubated at 37 °C overnight with 900 rpm agitation on a 1.5 mm orbit. Following overnight growth, cells were diluted 1:100 into fresh culture medium supplemented with antibiotic(s) in a 96-well deep-well plate and incubated for two hours, after which cultures were induced by the addition of ligands. Ligands were prepared as 1 molar stock solutions in DMSO. Cultures were incubated for an additional four hours and then harvested by centrifugation at 3500 x *g* for 10 minutes. Cell pellets were resuspended in 1 mL PBS pH 6.8 and 100 µL of cell suspension was transferred to a 96-well microtiter plate. Fluorescence assays were conducted using a multimode plate reader (Tecan M200 Pro). Absorbance was measured at 600 nm and fluorescence was measured using an λ_ex_ 550 nm and λ_em_ 610 nm.

Band stop variants were assayed using the workflow described above. To assay inverter variants, cultures were induced immediately after dilution into fresh medium supplemented with ligand and antibiotic(s) and incubated for a total of six hours, omitting the two-hour pre-induction growth step.

#### Statistical analysis and data visualization

Unless otherwise indicated, all data for fluorescence assays were collected from a minimum of three biological replicates and three technical replicates. Error bars represent the standard deviation of the mean of the biological replicates.

For the probability-of-inversion for each tested variant in the GROQ-Seq DMS assay, we used the results from the Bayesian inference with the Hill equation model to calculate the posterior probability that the transcriptional signal/output is lower at saturating concentrations of ligand than at zero ligand.

## Results and Discussion

We selected the TetR-family transcriptional repressor CymR from *Pseudomonas putida* as our candidate for mutagenesis for three reasons: (i) it has a well-characterized operator sequence, (ii) for a model biosensor it is relatively ‘information poor’ with limited prior mutagenesis and no crystal structure, (iii) and its native ligand cumate, a degradation product of p-cymene, retains some structural elements of cyclic monoterpenes, an industrially relevant class of molecules which lack good transcriptional biosensors^15^ (**Figure S1**). In our GROQ-Seq workflow, we situate all elements of a genetic circuit on a single plasmid, which allows for long-read DNA sequencing to identify library members with off-target mutations in the plasmid backbone that can later be excluded from analysis (**Figure 1a-1b**). We used a variant of CymR, CymR^AM^, as the template for the library. This variant contains the mutations S110G and A171V that have previously been reported to improve the dynamic range and reduce antagonism between sodium salicylate and cumate^21^. The *cymR^AM^* gene is expressed constitutively from a weak promoter paired with a bi-cistronic design translational control element (BCD)^34,35^. The function of CymR variants is linked to the expression of two reporter genes, *tetA* encoding a tetracycline efflux pump and the monomeric red fluorescent protein mScarlet-I. During GROQ-Seq, expression of the antibiotic resistance marker is linked to cell growth and fitness. As the TetA efflux pump does not degrade tetracycline, it is expected to yield a more consistent selective pressure over time when compared to other antibiotic resistance markers, such as beta-lactamases. The two genes are expressed as a bicistron under the control of the P*_cymRC_* promoter. To ensure quantitative measurements of function in the GROQ-Seq method, the expression level of TetA required tuning to produce a graded recovery of fitness across the range of expression levels achievable by CymR and its cognate promoter.

**Figure 1:**
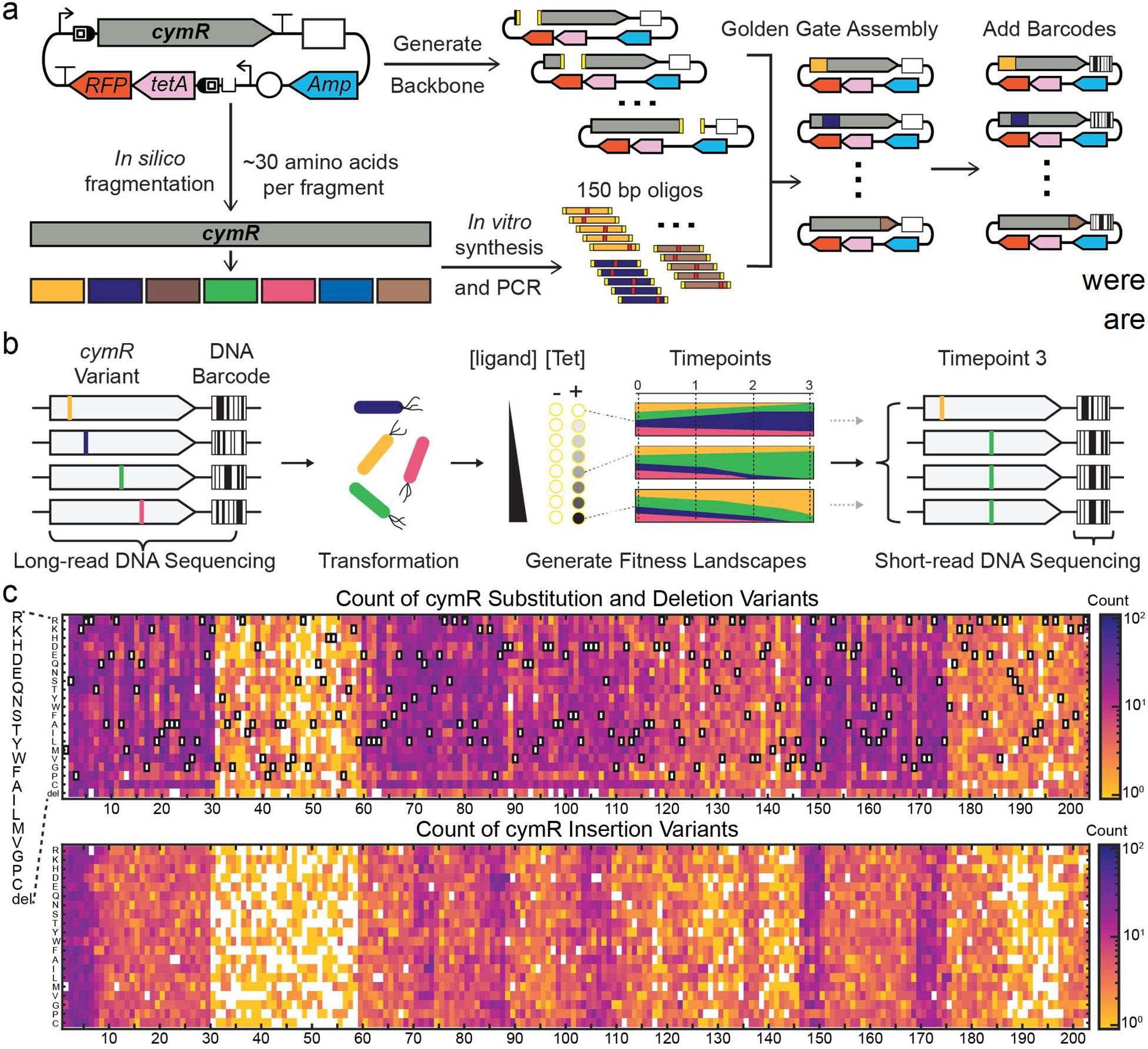
Workflow for construction and screening of a CymR deep mutational scanning library. (a) A plasmid backbone containing a reporter cassette, constitutive cymRAM repressor, and barcode site was used to generate the CymR deep mutational scanning library. The CymR sequence was divided into seven fragments in silico. Fragment-specific backbones and sub-pools were amplified via PCR, assembled by seven parallel BsaI Golden Gate reactions, pooled, and transformed into cells. Unique 18-bp DNA barcodes were added downstream of CymR to enable high-throughput tracking of variants. (b) The barcoded CymR library was assayed using a growth-based selection platform in 96-well plates under multiple ligand and concentration conditions. Long read sequencing was performed before any selection occurred, and short read sequencing was performed for each condition after the selection pressure was applied. (c) Heatmaps show the abundance of substitutions, insertions, and deletions across the CymR sequence in the library that had enough data generated in GROQ-Seq to generate functions.

To tune the circuit appropriately, we constructed a series of plasmids with different strength BCDs controlling the expression of TetA. Low levels of TetA expression can rapidly confer tetracycline resistance, and the P*_cymRC_* promoter has strong basal activity. Under these conditions, we determined that plasmids encoding an extremely weak BCD translational control element resulted in the best fitness gradient (**Figure S2**). We constructed the CymR DMS library using seven individual 150 bp fragments amplified from an oligonucleotide pool, each spanning 29 amino acids, and encoding all single amino acid substitutions, insertions, and deletions. This design encompasses approximately 40 mutations per residue (19 substitutions, 20 insertions, one deletion) for a total of 7887 unique sequences. To generate the *cymR* gene fragments, we used a derivative of the Saturated Programmable Insertion Engineering (SPINE) oligonucleotide design tool, which we adapted to perform DMS and reduce library redundancy. Briefly, the SPINE tool fragments the gene of interest into oligonucleotides of user-defined length. Each oligo encodes a single mutant gene fragment flanked by type II restriction sites, along with additional buffer sequences to normalize length and amplification tags at the termini^36^. We extended this tool to support DMS, including insertions and deletions and added an additional quality control step to ensure that no new Type II restriction sites unintentionally introduced via the mutations. To further reduce library bias, a deduplication step was incorporated into the code to eliminate redundant mutations caused by consecutive identical amino acids (e.g. the sequence VDDDA) where an insertion or deletion of an aspartic acid (D) would generate repetitive designs. This code supports AarI/PaqCI, a commonly used Type II restriction enzyme for library assembly, in addition to BsaI used by SPINE. The modified SPINE code is freely available on Github (Thyerlab/SPINE_Thyer). We note that another recently developed variation on the SPINE tool, DIMPLE, generates similar oligo designs and includes a graphical user interface for ease of use^24^. Oligonucleotide sub-pools and their corresponding plasmid backbones were amplified by PCR and ligated using Golden Gate assembly, after which DNA barcodes were appended to each variant in the library by PCR and re-circularized.

We transformed the barcoded CymR DMS library into *E. coli* cells, bottlenecked the population of transformants to an estimated 200,000 CFUs, and sequenced the resulting bottlenecked library with long-read (Oxford Nanopore) sequencing to determine the DNA barcode and the CymR coding sequence for each variant in the library population. The long-read sequencing results indicated that the library contained more than 95% of all possible substitutions (Figure 1c). Variations in abundance of the different CymR fragments were observed with the majority of the missing variants found in fragment two (spanning residues 31 to 59), which had a significantly lower abundance than the other fragments of CymR. The library contained more than 85% of all possible insertion mutations, and more than 82% of the possible deletion mutations. Insertions and deletions showed lower abundance across all fragments compared to substitution mutations, which has been previously reported with similar workflows, and it is unclear what led to this imbalance given all mutation types are encoded within the same subpools^24^. The variations in abundance between the CymR fragments in the library are most likely due to different assembly efficiencies of the seven sub-libraries, which were then pooled prior to DNA purification and transformation. We predict that maintaining separate sub-libraries through the assembly and barcoding reactions and only pooling supercoiled plasmid DNA prior to the final transformation will minimize this issue^24^.

To characterize the effect of mutations on the CymR sensor phenotype, we measured *in vivo* dose-response curves of each CymR variant in the bottlenecked library using a quantitative growth-based method similar to the approach described by Tack et al^29^. Briefly, we used an automated, growth-based protocol to measure the growth rate of cells expressing different CymR variants in 24 different sample conditions: with and without tetracycline and over a range of concentrations of three ligands: perillic acid (PA), perillyl alcohol (Per-OH), and S-limonene (S-Lim). We used DNA barcode sequencing to measure the growth rate associated with each CymR variant in a pooled assay format for each of the 24 samples. Finally, we used a well-characterized set of CymR variants as a “protein function calibration ladder” to convert the growth rate data into dose-response curves of the CymR circuit as a function of PA, Per-OH, and S-Lim concentrations (Figure S3).

Analysis of the results indicated that our GROQ-Seq experiment was able to capture a comprehensive phenotypic landscape including multiple non-standard phenotypes using PA, however, the results with Per-OH and S-Lim indicated that no single amino acid mutations were capable of generating a CymR variant that could significantly respond to either of these ligands. A small number of variants that demonstrated a slight response to these ligands but displayed high error measurements across replicates were reconstructed and evaluated using clonal plasmids. These were found to have no significant response to Per-OH and S-Lim, and all subsequent analysis was performed on the PA population (Supplementary File 1, Table S3).

Several parameters of the dose-response curves for an allosteric transcriptional regulator can be analyzed to assess the phenotypic impact of each mutation to CymR, collectively yielding a broad phenotypic landscape. One key parameter is the level of transcription when no ligand is present, effectively representing the off-state of the circuit or basal level of transcription. Wildtype CymR binds tightly to the DNA, resulting in a very low level of transcription when no ligand is present. We calculated the predicted transcriptional activity for all variants in the library in the absence of ligand as the function G_0_ (Figure 2a). Single amino acid substitutions show a distinct banding pattern, where specific residues are generally amenable to mutation without impacting function, while other positions have an extremely low tolerance for any amino acids besides the wildtype residue. In contrast, at a majority of positions in CymR, any single amino acid insertion or deletion results in a large deleterious impact to protein function, effectively abolishing transcriptional repression. CymR only tolerates insertions or deletions at the termini of the protein, in unstructured loops in between alpha helices, and in residues 169-187 in the two C-terminal alpha helices which form the dimerization interface.

**Figure 2:**
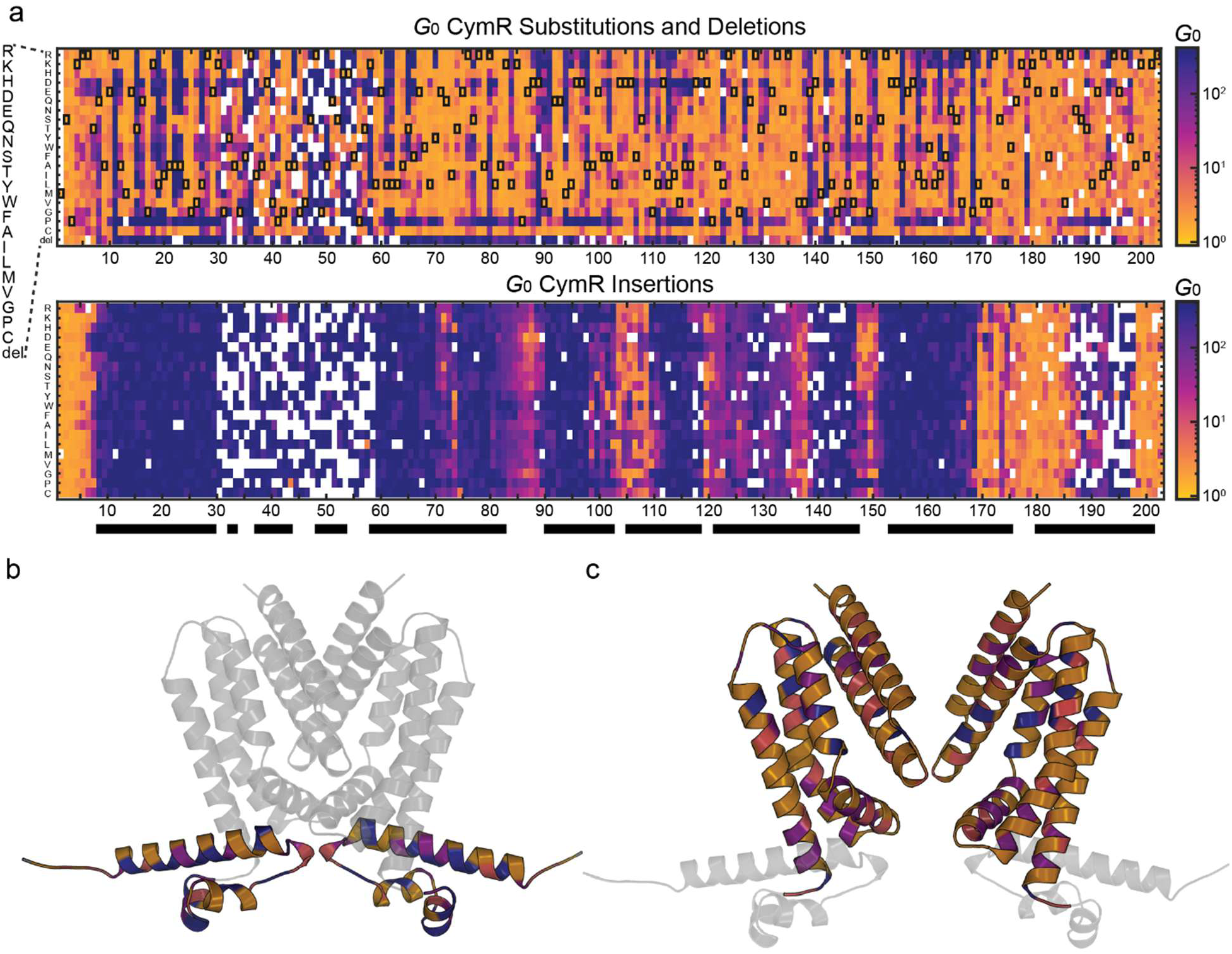
Analysis of the G0 score for all tested variants of CymR. (a) Heatmaps showing the G0 score for all tested variants. Black boxes in the substitution and deletion heatmap indicate WT residues. White spaces indicate a variant that was not present in the library. Black bars along the bottom indicate the alpha helices of the CymR structure. Alphafold-generated CymR structures shown (b) with the DNA binding domain highlighted or (c) with the Ligand binding domain highlighted and the two dimers separated slightly to avoid overlapping. The color scale indicates the percentage of tested variants at each residue which raised the G0 by more than two standard deviations above the wildtype. Blue indicates more than 80% of variants, purple indicates between 60-80% of variants, pink indicates between 40-60% of variants, and orange indicates between 0-40% of variants.

We calculated the percentage of single amino acid substitutions at each position which resulted in a G_0_ score more than two standard deviations above the wildtype CymR when no ligand is present and mapped these scores onto an Alphafold-generated structure of CymR, for both the DNA binding domain and the ligand binding domain (Figure 2b-c). In the DNA binding domain, mutations which generally tolerate many amino acid mutations are oriented away from the DNA binding interface of the helix-turn-helix domain, whereas residues which are predicted to be DNA-facing are substantially less tolerant of mutations (**Figure2b**). This distribution accounts for the striking banding pattern observable in the N-terminal region of the G_0_ heatmap and indicates that substitutions in the DNA binding domain which elevate the G_0_ score adversely impact the ability of the DNA binding domain to effectively bind to the P_cymRC_ promoter sequence.

Similarly, in the ligand binding domain, most residues which orient the amino acid side chain outwards to the exterior solvent exposed surfaces of the protein are highly tolerant of substitutions, while residues with side chains either within or adjacent to the internal ligand binding cavity are much less permissive (**Figure2c**). In particular, residues 98, 102, 139, 142, 146, and 156 all directly extend into the hydrophobic core of the protein, and each is highly intolerant of mutation, suggesting that these residues are critical for correct folding of CymR. Additionally, residues 165 and 169 which occupy the center of the dimerization interface are similarly immutable, where all possible substitutions elevated the G_0_ score. Finally, across most of the CymR sequence, substitution with a proline residue elevated the G_0_ score, which is not unexpected due to the disruption of local protein structure caused by the constrained cyclic backbone.

A second parameter for ligand-inducible transcriptional regulators is the induction fold-change, or the ratio of transcriptional signal between the maximum ligand concentration and when no ligand is present. We calculated the fold-change between the maximum concentration of PA tested, 2500 μmol/L, and the previously described G_0_ score, and mapped this from 10^-2^-10^3^ (Figure 3a) Here, values greater than one (blue) indicate CymR mutants with higher transcriptional signal when induced with 2500 μmol/L PA, with larger values indicating a wider difference between the zero and high ligand states. The wildtype CymR^AM^ has a high fold-change score of 180, and is shown as a dark blue color inside of black boxes, indicative of good sensitivity to the ligand. Values close to one indicate minimal change between 2500 μmol/L PA and 0 μmol/L PA and are shown with a pale turquoise color. This phenotype indicates that the protein is no longer exhibiting allosteric regulation, where it may not repress transcription at all, or may not de-repress in the presence of ligand. Interestingly, we also identified a large number of variants with a calculated fold-change of less than 1, designated with a red color. These variants show high fitness in the absence of PA, and reduced fitness at high concentrations of PA, effectively displaying an inverted response compared to the wildtype protein.

**Figure 3:**
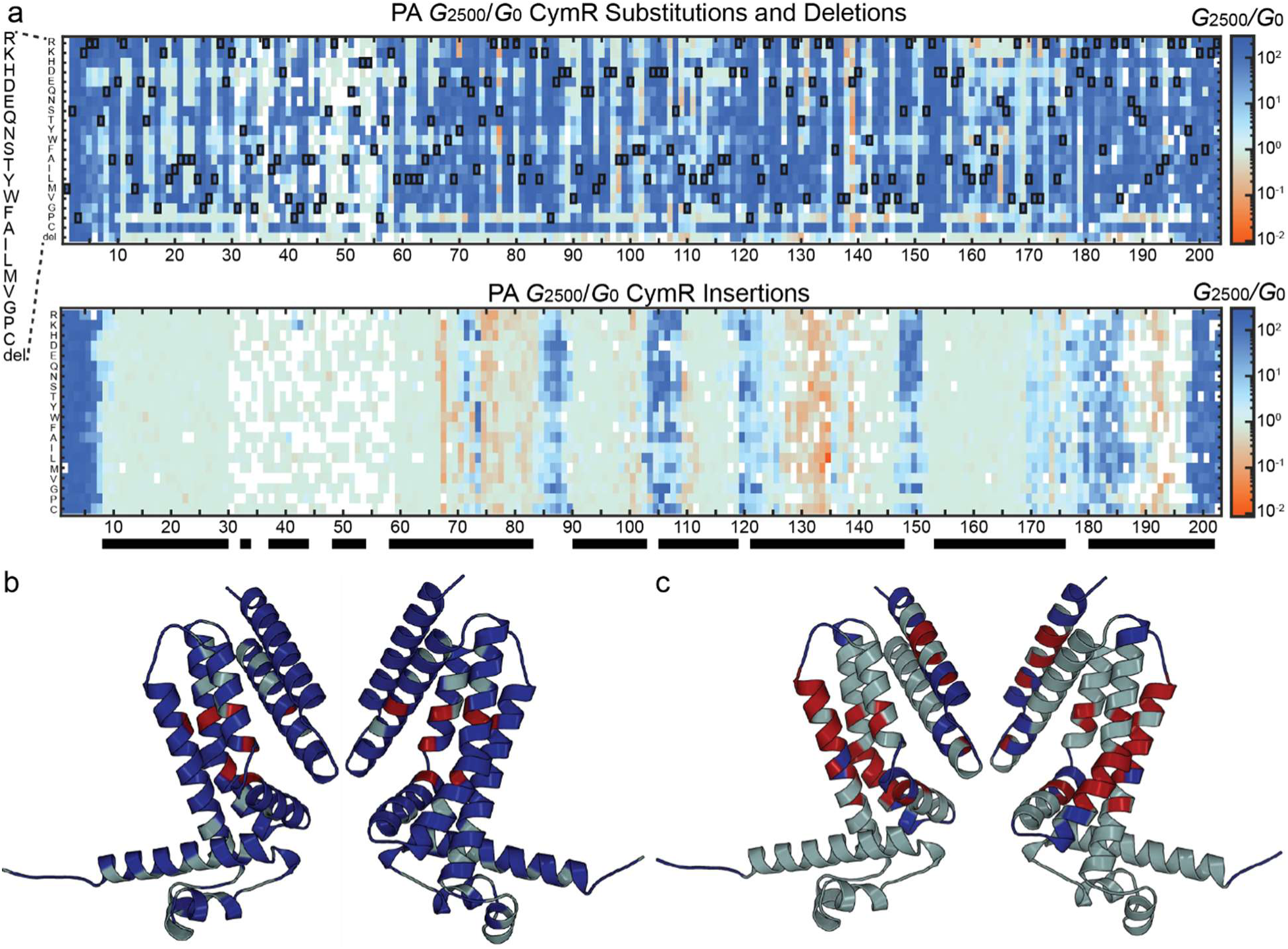
Analysis of the fold change for every tested variant of CymRAM. (a) Heatmaps showing the transcription rate at 2500 µmol/L PA divided by the transcription rate at 0 µmol/L PA (G2500/G0) CymRAM variant Black boxes in the substitution and deletion heatmap indicate WT residues. White spaces indicate a variant that was not present in the library. Black bars along the bottom indicate the alpha helices of the CymR structure. CymR structures separated slightly to show both dimers without overlapping for (b) substitutions and (c) insertions with each residue colored to indicate the likelihood that mutations at that residue lead to an inverted phenotype. Red indicates residues where more than 20% of the tested variants had a greater than 97.5% confidence of being an inverted phenotype. Blue indicates residues where less than 20% of tested variants had an inverted phenotype and mutations were more likely to be tolerated than not. A grey color indicates residues where mutations were more likely to lead to a no-fold change variant rather than a high fold-change variant.

This dataset reveals important information about CymR^AM^. First, while most variants that exhibit a fold change close to one also exhibit an elevated G0, a small number of variants possess low G_0_ scores along with a fold-change close to one. These would indicate variants which maintain tight binding to the P*_cymRC_* promoter, but no longer either bind the ligand or change conformation upon ligand binding to release from the DNA when 2500 μmol/L PA is present. We have identified 17 positions where more than 20% of substitutions resulted in this phenotype and marked their location in the CymR structure (**Figure S4**). Eight of these positions (132, 161, 168, 170, 173, 177, 182, and 186) appear to have side chains near the opening to the central cavity, and mutations at these locations may result in occlusion of the ligand binding pocket. A further five positions (74, 100, 101, 103, and 110) have side chains directly lining the binding pocket, which are potentially required for ligand recognition. Finally, while four of these positions (39, 68, 134, and 141) do not have side chains that are likely to form direct interactions with the ligand, they may form second shell interactions to position other residues or facilitate signal transduction between the ligand binding domain and the DNA binding domain. Residue 39 is the only position among this set which falls within the DNA binding domain, and position 68 has a large side chain oriented directly above the first helix of CymR in the DNA binding domain, and we hypothesize that these two positions are the most likely to play a role in signal transduction.

Most single amino acid insertions within structured alpha helices of CymR result in a fold-change of close to one, indicating variants that cannot bind to the DNA or respond to PA. This result is consistent with previous reports that have shown insertions to be well tolerated in unstructured loops and at the termini of a protein, while being deleterious in more structured parts of the protein^37–39^. This data provides good evidence that the phenotypic landscape measurement is broadly capturing the expected biophysical behavior from a population of ligand-inducible transcriptional repressor variants. In addition to their generally deleterious impact on CymR function, the phenotypic landscape reveals that inversion-of-function mutants are not rare, and while they can be found in the population of amino acid substitutions, it is insertions that are a key driver of this phenotype. We used the GROQ-Seq data and associated uncertainty across all tested concentrations of PA to calculate the probability-of-inversion for each tested variant (Methods). At least one mutation with a greater than 97.5% probability of conferring inversion-of-function was observed in more than 25% of all residues in CymR^AM^ and comprised more than 10% of all insertion variants. To investigate inversion of function “hot-spots”, residues where more than 20% of mutations result in an inverted phenotype have been mapped onto the model of CymR (Figure 3b-c) In contrast to the insertion mutants, that preserved the normal biosensor response and predominantly occur in flexible loops or near the termini, those conferring an inverted phenotype cluster within a series of discrete hotspots, distributed in both the sequence and structure of CymR, but largely falling in the center of alpha helices. Substitution mutations which resulted in an inverted phenotype were far less common, however, position 139 was an exception, as thirteen of the nineteen possible substitution mutations at position 139 resulted in an inverted phenotype. E139G was previously reported to result in a CymR variant with an inverted phenotype and the GROQ-Seq results agree with this^20^.

A third parameter to evaluate is the ligand concentration that yields a fold-change halfway between the off-state and maximal induction (EC_50_) which can be a measure of sensitivity to a given ligand^36–38^. However, the EC_50_ is also influenced by other factors within a system, such as the concentration of the transcription factor^40^. Thus, mutations that decrease CymR protein production, perhaps via less efficient translation at the N-terminus or decreased stability and half-life, may also have a lower EC_50_. We calculated and mapped the EC_50_ for all CymR variants with PA onto the CymR structure (**Figure S5**). A number of substitution mutations across CymR were found to lower the EC_50_. However, some of these mutations also had elevated G_0_ scores, which may complicate the evaluation of the EC_50_. Additionally, a large number of insertion mutations that were calculated to have a fold-change of less than one, indicating an inverted phenotype, also showed lower EC_50_ than the wildtype CymR^AM^ After analyzing the Groq-Seq dataset we sought to empirically validate several of the observed phenotypes. To identify a subset of CymR variants with broadly improved ligand response, scores from the Hill function fit for PA were ranked by fold-change (G_2500_ / G_0_) to identify CymR variants with higher maximum induction and/or tighter off-states. We primarily selected variants for characterization based on the largest differences between the on- and off-states, with additional filtering to sample diverse positions in the sequence, and, where possible, low measurement error and higher independent barcode counts. We constructed a subset of these variants with low apparent off-states for characterization and assayed them in a two-plasmid context that fully separated the CymR variants from the fluorescent reporter. To confirm that these variants had high sensitivity for PA, we induced the variants with 50 µmol/L PA and compared them against the wildtype CymR^AM^ (**Figure S6**). From this subset, the three mutations at spatially separated positions with the highest induction at 50 µmol/L PA as compared to wildtype CymR^AM^, G25D, V137T, and I91F, were chosen for further interrogation (Figure 4a-b).

**Figure 4:**
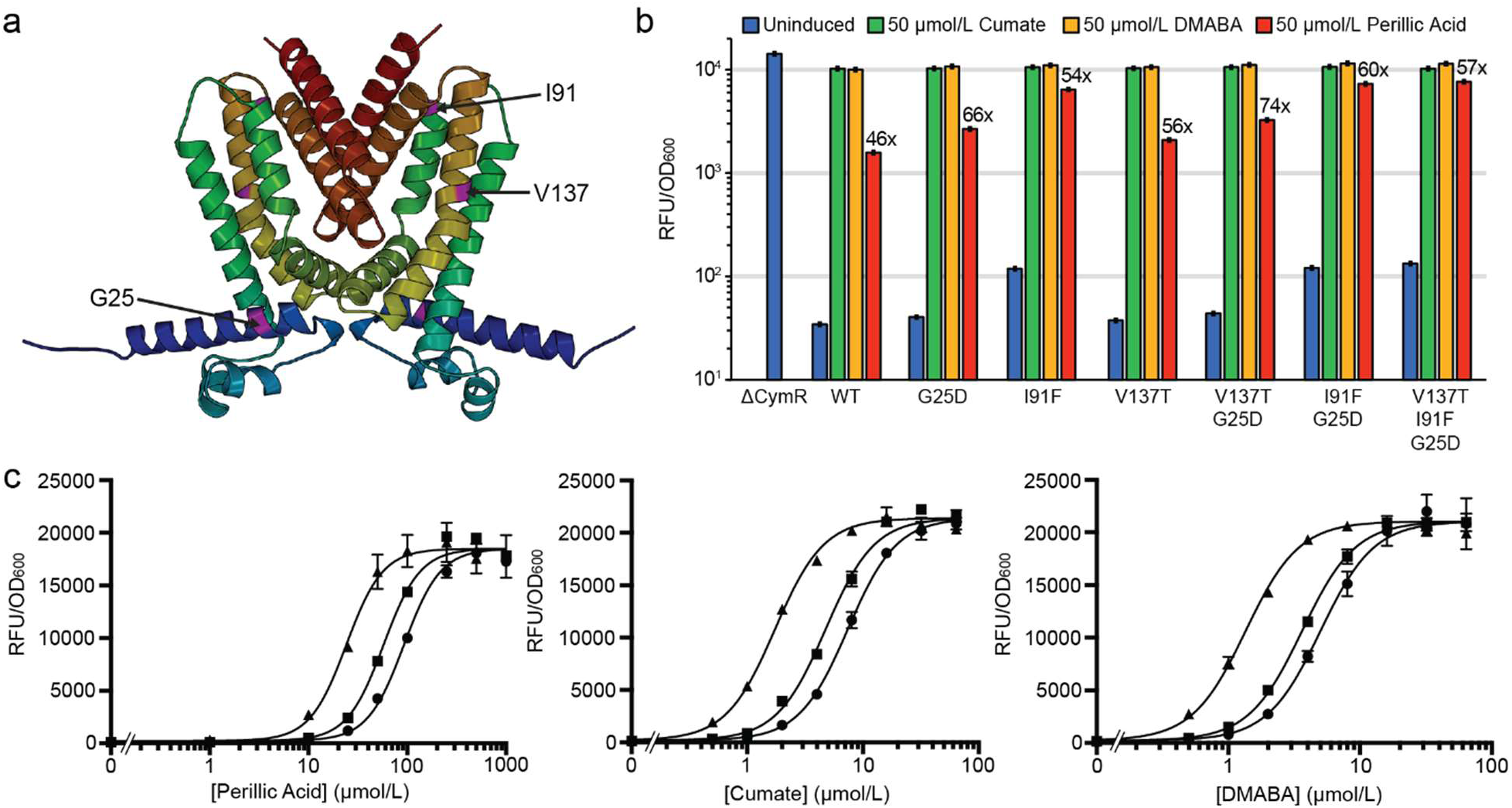
Variants with improved response to perillic acid. (a) Structural representation of CymR highlighting sites tested for improved perillic acid response. (b) Fluorescence assays comparing CymR^AM^, three single mutants (G25D, I91F, and V137T), and their combinations in response to 50 µmol/L perillic acid, shown on a log10 scale. (c) Dose-response curves for Perillic Acid, Cumate, and DMABA, comparing CymR^AM^, the double mutant G25D + V137T, and the triple mutant G25D + V137T + I91F.

The G25D mutation is found in the N-terminal DNA binding domain and confers a slight increase in sensitivity to PA while maintaining the low basal output for CymR^AM^. V137T also provides a moderate increase in induction with submaximal concentrations of PA with a minimal increase in off-state signal, while I91F confers a large increase in sensitivity but with a concomitant increase in the basal signal. Given the three mutations occur in different helices and have no direct interactions, we hypothesized that the observed effects may be additive and evaluated several combinations. Consistent with this hypothesis, we found that the combination of G25D and V137T maintained a low basal signal and yielded the highest fold-change induction in response to 50 μmol/L PA. All combinations that included I91F show an elevated basal signal alongside an improved induction at 50 µmol/L PA (Figure 4b). We hypothesized that it may be possible to lower the basal signal by increasing the amount of CymR expressed, while preserving the increased ligand sensitivity. However, while titrating CymR expression using a series of bicistronic design ribosome binding sites (BCDs) was able to reduce the basal signal, maximal induction of the circuit was simultaneously decreased, resulting in minimal overall improvement to the fold change (**Figure S7**).

To evaluate the effect of these mutations across different ligands, we compared the induction of CymR^AM^, the double mutant G25D and V137T, and the triple mutant containing I91F using PA and the canonical ligands cumate and 4-(Dimethylamino)benzoic acid (p-DMABA) (Figure 4c). Across all three ligands, both the double mutant and triple mutant displayed a lower EC_50_ than CymR^AM^, with the inclusion of I91F resulting in a more than 3-fold improvement in sensitivity to all three ligands. Based on these results, we conclude that the double mutant represents an improved, more sensitive version of the CymR^AM^ sensor for use in genetic circuits that maintains the desirable low basal signal, while the I91F mutant, either alone or in combination, may prove useful as a starting point for further efforts to broaden the substrate scope of CymR towards additional ligands for which it has low sensitivity.

Transcriptional repressors may demonstrate vastly different performance under different assay conditions. To further inform the use of CymR as a component of genetic circuits, we assessed induction of CymR^AM^, the double mutant containing G25D and V137T, and the triple mutant containing I91F during culture in five different bacterial media; M9 salts supplemented with glucose and casamino acids, M9 salts supplemented with glycerol and yeast extract, lysogeny broth (LB), 2xYT, and terrific broth (TB) (**Figure S8**). While genetic circuits were functional under all media conditions, more complex media resulted in reduced fold-change across all three variants, and surprisingly reduced fluorescence signal. Fluorescent background from cells lacking the mScarlet-I reporter used for normalization was uniformly low, and repressor off-states were not significantly impacted, with the changes in fold-induction driven primarily by reduced reporter expression.

Although the CymR phenotypic landscape indicated that inversion-of-function mutants were widely distributed throughout the sequence, analysis of the data indicated that while most were fully de-repressed in the absence of ligand, they were incapable of complete repression, even at high concentrations of PA (**Figure S9**). For characterization, we selected a subset of mutations that conferred a phenotype with low to moderate signal in the absence of PA, and tighter repression at high concentrations, a phenotype likely to result in a desirable high fold-change. We constructed 12 putative inverted variants, prioritizing those which showed strong repression at high levels of PA and were widely distributed across the sequence of CymR, of which 11 yielded observable repression at 50 µmol/L PA (**Figure S10**). We selected six of these variants, substitutions S77M and V145P, and four insertions within an inversion hotspot spanning residues 131-139, for further characterization (Figure 5a-b). In addition, we included a previously reported inversion mutant, E139G. We induced the selected variants with both 50 µmol/L and 1 mmol/L PA, and all were found to be more sensitive than E139G, with two variants (R133_N134insC and V138_E139insL) capable of >10-fold change in fluorescence. While E139G did not demonstrate repression at 50 µmol/L PA, it tightly repressed expression at 1 mmol/L (Figure 5b).

**Figure 5:**
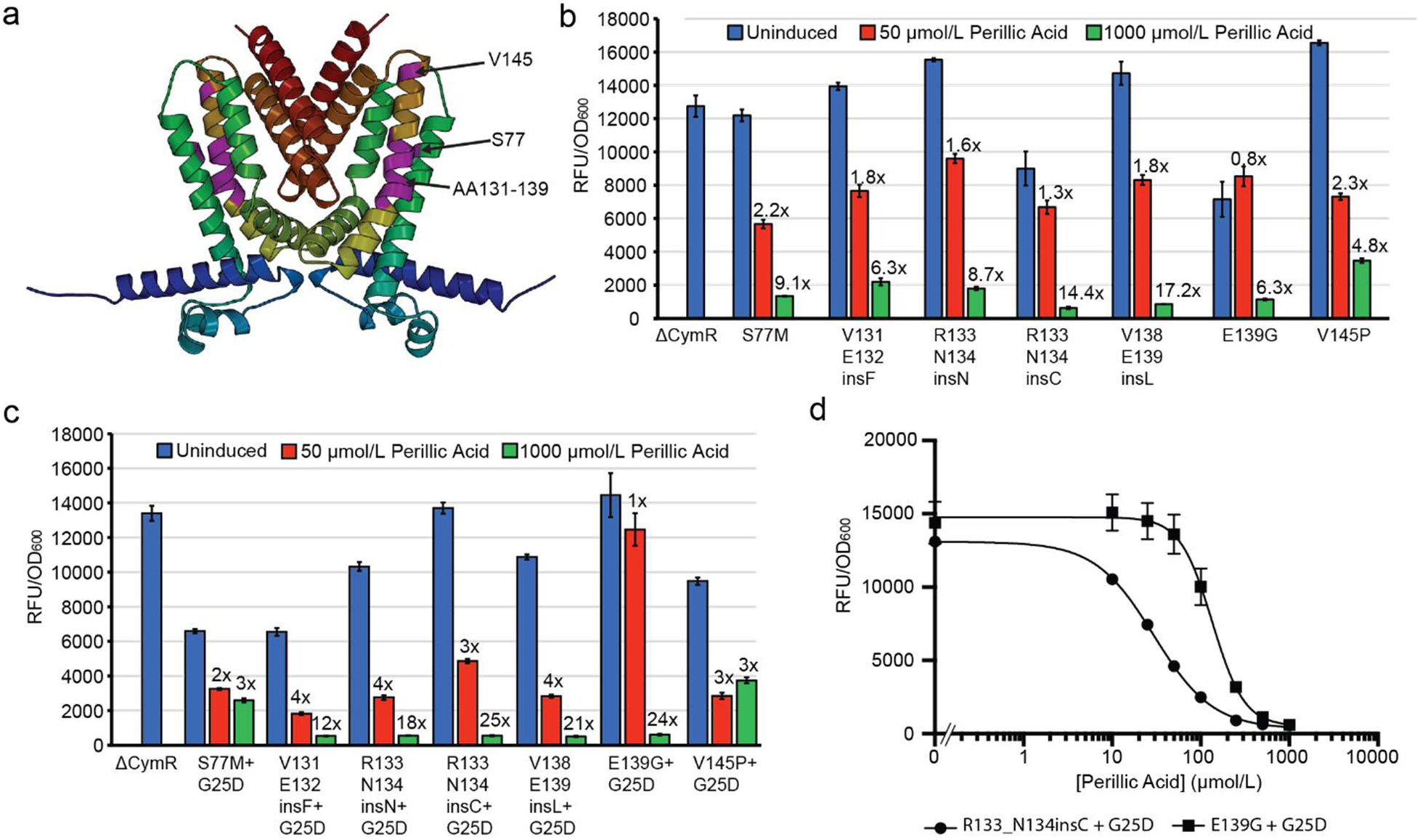
Mutations conferring inverted phenotypes in CymR. (a) Structural representation of CymR highlighting sites associated with inverted phenotypes. (b) Fluorescence assays with 50 µmol/L and 1 mmol/L perillic acid comparing inverted variants from this study with the previously reported E139G variant (c) Fluorescence assays testing inverted variants combined with the G25D mutation to evaluate effects on DNA binding and repression strength. (d) Dose response curves for perillic acid comparing the R133N134insC + G25D inverter variant with E139G + G25D.

We hypothesized that the G25D mutation located in the DNA binding domain which improved induction with PA may enhance the performance of the inversion-of-function mutants. In most cases, addition of G25D increased repression in the off-state and improved the overall fold-change in fluorescence (Figure 5c) with the best performing variants exceeding a 20-fold change in fluorescence. Interestingly, all mutations within the 131-139 region were enhanced by G25D, while the two characterized inversion-of-functions variants outside of this region, S77M and V145P, were not. Considering the improvement in PA sensitivity observed when pairing G25D and V137T, these two regions appear to yield a variety of additive or synergistic phenotypes and may be good candidates for co-mutagenesis. The sensitivity of two of the best performing variants, R133_N134insC + G25D and E139G + G25D, was determined with PA, and the two canonical ligands cumate and DMABA (Figure 5d**, Fig S11**). Both mutants retained the inversion-of-function phenotype across all three ligands, with R133_insC_N134 + G25D possessing a lower EC_50_, particularly in response to PA. Collectively, our results significantly expand both the number and performance of inversion-of-function mutants in CymR.

An advantage of the GROQ-Seq methodology is the generation of a full dose-response curve for each variant in the library, which can reveal novel phenotypes. One example is a band-stop phenotype, where repression occurs only at intermediate ligand concentrations. The band-stop phenotype is inconsistent with the monotonic, sigmoidal form typically used to analyze transcription-factor dose-response curves (e.g., the Hill equation). To detect possible band-stop phenotypes, we used results from a Gaussian Process (GP) model applied to the GROQ-Seq PA dataset. Specifically, we searched the model outputs to identify variants with a dose-response that was significantly negative at low ligand concentration and significantly positive at a higher ligand concentration, of which 34 mutants met these criterion (**Table S4**).

From this list, we constructed a single variant, Y70_E71insN, and confirmed that it yielded a band-stop phenotype with maximal repression observed at 50 µmol/L PA and an approximate bandwidth of 100 µmol/L (Figure 6a-b). To our knowledge, this is the first reported occurrence of a band-stop phenotype in a transcriptional repressor conferred by a single mutation^29^. In agreement with the previous results, addition of G25D to Y70_E71insN increased sensitivity to PA and yielded maximal repression at 25 µmol/L PA, while also increasing induction at 1 mmol/L PA. However, the increased sensitivity was accompanied by reduced signal in the absence of ligand and thus degraded the overall band-stop phenotype. To test the sensitivity of the band-stop phenotype to changes in biosensor concentration we titrated CymR^AM^ Y70_E71insN expression using a gradient of stronger BCD translational control elements (Figure 6c). With increasing protein expression, greater operator site occupancy results in increased repression across all concentrations of ligand, with band-stop behavior diminishing beyond the 1.5K BCD.

**Figure 6:**
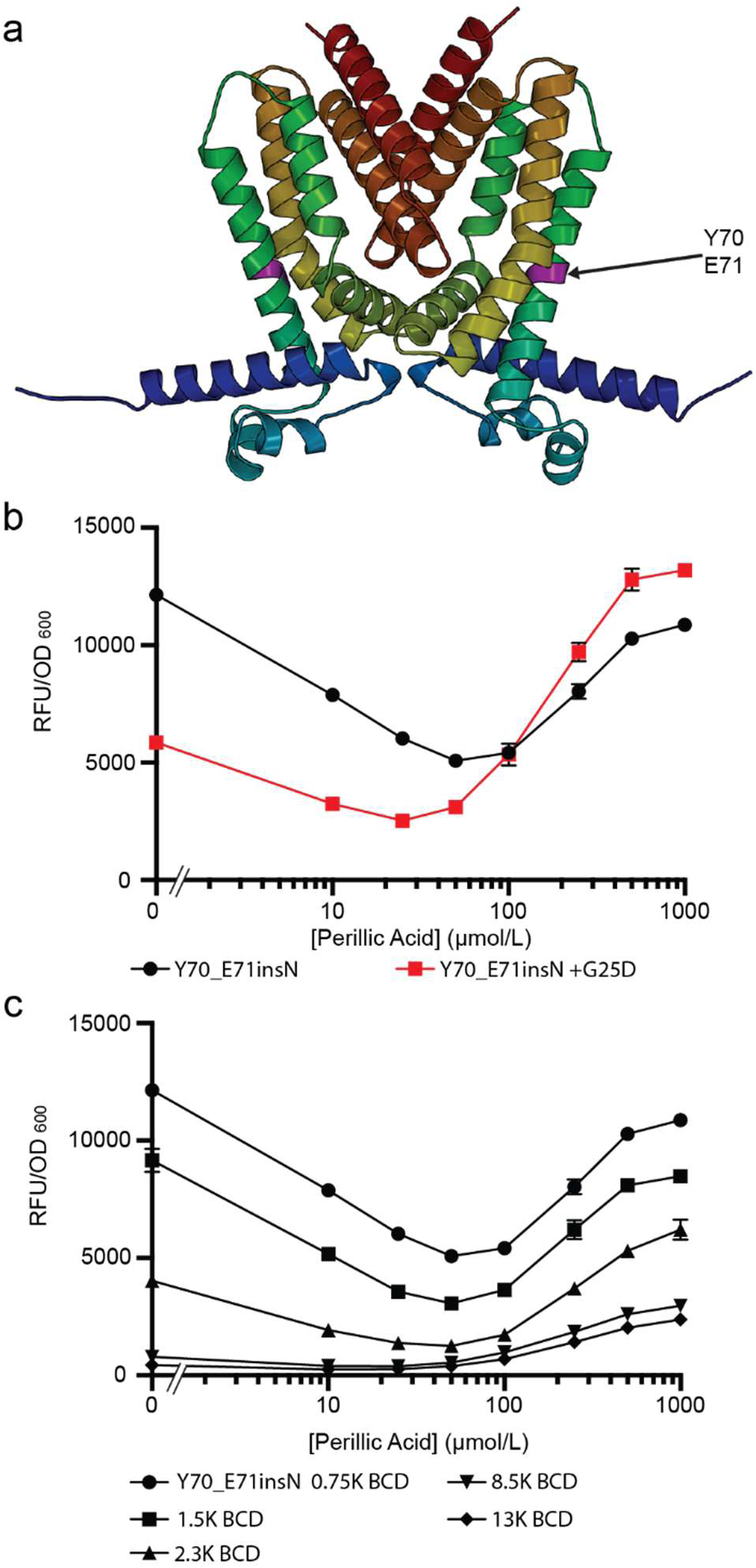
The Y70_E71insN mutation confers a novel band-stop phenotype in CymR^AM^. (a) Structural representation of CymR^AM^ highlighting the Y70_E71 insertion site associated with a band-stop phenotype. (b) Fluorescence assays showing the Y70_E71insN variant exhibits a band-stop phenotype in response to perillic acid, which is accentuated when combined with the DNA-binding mutation G25D. (c) Fluorescence assays testing the effect of increased CymR^AM^ expression using stronger BCD translational control elements.

## Conclusion

Here we combine DMS aided by computational generation of the mutagenic oligonucleotide fragments and GROQ-Seq to thoroughly interrogate the mutational landscape of the ligand-dependent transcriptional repressor CymR. A derivative of the SPINE oligonucleotide design tool was developed that enables a simple workflow for generating DMS libraries of any size from oligonucleotide pools. The CymR library comprised over 7800 unique single mutations which were screened using GROQ-Seq across 24 different ligand and antibiotic concentrations. From the resulting phenotypic landscape, we identified mutations conferring a variety of expected and novel phenotypes, including improved repression and ligand sensitivity, inversion-of-function, and the novel band-stop phenotype. Additionally, we identified residues in CymR which may contribute to a variety of protein functions. We empirically validated multiple inversion-of-function variants spanning several loci in CymR, and we further engineered them to improve off-state repression. These mutants represent a family of new regulators that are seamlessly compatible with genetic circuits using P*_cymRC_* promoter or the isolated cymO operator.

Previous characterization of reverse-TetR variants indicated that the mutations resulted in a more disordered and less structurally stable protein that was unable to bind DNA in the absence of ligand. The ligand binding both stabilized the protein and allowed it to adopt a configuration that binds the DNA operator and represses transcription. Our results indicate that a similar inverted phenotype is achievable in CymR by single amino acid substitutions at a limited number of locations, and far more broadly in the sequence through single amino acid insertions. It seems likely that these mutations are having a similar impact on CymR, although the occurrence of inversion-of-function mutations in both the ligand binding helices and the dimerization interface suggests different mechanisms to impact DNA binding.

Cryptic phenotypes such as inversion-of-function or a band-stop are rarely observed in mutagenesis studies of transcription factors, often due to filtering steps that explicitly eliminate variants that fail to exhibit the expected behavior of a functional transcriptional repressor. In contrast, GROQ-Seq measures the entire phenotypic landscape and samples a variety of induction profiles rather than a handful of extreme ligand concentrations and was highly successful in capturing these non-standard phenotypes. Not only were these phenotypes observed, but near-complete mapping of the phenotypic landscape revealed that inversion-of-function mutants were surprisingly abundant, representing more than 3% of substitutions and 10% of insertions. This indicates that insertions, a class of mutation rarely sampled by traditional methods of mutagenesis, are highly enabling in certain contexts and can facilitate the acquisition of phenotypes that would be difficult to access through single nucleotide changes. Furthermore, although inversion-of-function hotspots were clearly apparent, individual mutations conferring this phenotype were widely distributed in both sequence and structure, suggesting several biophysical routes to refactor the conformational changes that drive allosteric signal transduction. While this phenomenon has not been extensively investigated, inversion-of-function hotspots do not appear to be conserved across TetR-family transcriptional repressors, which suggests multiple mechanisms by which this phenotype can arise^41–43^.

It is unclear why no CymR variants with significantly improved recognition of the alternate ligands limonene and perillyl alcohol, less oxidized derivatives of perillic acid, were identified. Possible explanations include the requirement for an aromatic acid and the inability of any single mutations to substantially rearrange the ligand binding pocket to accommodate alternate functional groups at this position, or the prior mutations acquired during construction of the CymR^AM^ scaffold favoring tighter interactions with canonical ligands. Interestingly, while the E139G mutation was previously described in the context of wild-type CymR, we evaluated it in both contexts and were only able to demonstrate a robust inversion-of-function phenotype in the CymR^AM^ background. The reasons for this discrepancy are not clear, but it does indicate that the S110G and A171V mutations in CymR^AM^, both of which lie within the ligand binding cavity, result in significant epistasis for additional mutations in the local environment, even without direct side chain interactions. The lack of variants with even a small response to the alternative ligands, along with the limited set of known ligands, suggests the CymR biosensor lacks the innate promiscuity of other well-characterized TetR-family repressors which are associated with multidrug efflux pumps and cellular response to xenobiotics^44^.

To guide future mutagenesis experiments using the GROQ-Seq workflow, we have identified three key changes to improve data collection and quality. First, we observed an imbalance in the ratio of each CymR sub-pool in our DMS library, and while overall library coverage was high, mutations within the second fragment were underrepresented. We hypothesize that this resulted from pooling the Golden-Gate assemblies of the sub-libraries together prior to transformation into cells. Instead, each sub-library could be assembled and barcoded individually while ensuring sufficient oversampling at each step, prior to a final pooling step. This workflow has been documented to reduce this bias and requires no significant changes to the library design^24^. Second, independent barcoding of the individual sub-libraries is expected to improve the quality of the data collected from GROQ-Seq by reducing barcode re-use. Both this work and prior GROQ-Seq investigation of LacI observed barcode re-use at much higher frequency than expected despite different barcode designs and barcoding strategies. While the basis for this remains unclear and the level of barcode re-use was not sufficient to impact data recovery, a parallel barcoding strategy represents a simple change to the workflow that also serves to representation of each mutagenic sub-library.

Finally, we propose that a key consideration for the design of future GROQ-Seq selection circuits should be to evaluate multiple selectable markers and identify the one with sufficient dynamic range and resolution to capture all circuit dynamics of interest. Here, the TetA tetracycline efflux pump was used to mediate cell fitness in response to tetracycline. However, even very low expression of TetA can result in full resistance and furthermore, high expression of TetA, a membrane protein, may confer a fitness burden. This results in a relatively steep change in cell fitness over a small gradient of CymR activity, providing finer resolution for small changes in protein function. Additional circuit designs may be considered that prioritize measuring fitness changes over the widest possible range of selection marker activity or expression. Examples may include stoichiometric binding proteins such as the *Sh*-ble protein or a far less active selectable marker, e.g. a catalytically impaired enzyme, that can be titrated over a wide expression gradient^45^.

The GROQ-Seq phenotypic landscape measurements represent a scalable approach to generating large protein sequence-to-function datasets. The sole requirement is for protein function to be connected to either expression or activity of a selectable marker, which can be achieved using creative design of genetic circuits for many protein families beyond transcriptional regulators. The collection of rich, high-resolution datasets of protein function will underpin new data-driven efforts in biological engineering and currently represents a limiting step towards developing better models for interrogating the interplay between protein sequence, structure, and function. Critically, these workflows capture numerous cryptic phenotypes that would not have been predicted nor observed from previous experiments, thus providing the most complete sampling of protein phenotypes possible to best inform our understanding of protein function, design, and engineering.

## Supporting information

cymr_variant_table

Plasmid Maps

## Abbreviations

DMS: deep mutational scanning
PA: perillic acid
GROQ-Seq: growth-based quantitative sequencing
DMABA: 4-(Dimethylamino)benzoic acid

## Author Contribution

Z.J., X.L., D.R., and R.T. conceived the study, designed the experiments, and coordinated the experimental work. Q.W. developed modifications to the SPINE code. Z.J., X.L., Q.W., D.L.K., N.A., O.V., A.R.G., and D.R. performed the experiments. Z.J., X.L., D.R., and R.T. analyzed the data. Z.J., X.L., D.R., and R.T. wrote the manuscript.

## Acknowledgements

This work was supported by funding from the National Institute of Standards and Technology (R.T., 70NANB21H102). We would like to thank the Genetic Design and Engineering Core at Rice University for their assistance with DNA sequencing.

## Supporting Information

Data used to analyze the GROQ-Seq results are provided in supplementary file 1. Additional experimental methods, figures S1-S11, Tables S1-S4, are provided in supplementary file 2. GenBank files containing annotated DNA sequences for plasmids used in this work are included. Supplementary Data are available at NAR Online

## Data Availability

Data used for the GROQ-Seq analysis is provided as a csv file, Supplementary file 1 Table S3 provides an explanation of the data in each column.

