## Supplementary Information for "Mapping the phenotypic landscape of a transcriptional repressor using Deep Mutational Scanning and Growth-based Quantitative Sequencing"

**Affiliations:**
[1] Systems, Synthetic, and Physical Biology, Rice University, Houston, TX
[2] Department of Chemical and Biomolecular Engineering, Rice University, Houston, TX
[3] Department of Bioengineering, Rice University, Houston, TX
[4] National Institute of Standards and Technology, Gaithersburg, MD, USA

This file contains:
Supplementary methods
Supplementary figures S1-S11
Supplementary Table S1-S4

**Supplementary methods**

<u>Selection of Normalization Controls and Protein Function Calibration Ladder for GROQ-Seq</u>

Two different types of reference, spike-in variants were used to calibrate the GROQ-Seq DMS data: firstly, normalization controls which were used to normalize the barcode read counts in each sample to account for sample-to-sample variability in DNA extraction efficiency and/or barcode PCRs, and secondly, a set of 16 CymR variants, with a range of different functional responses, which were used as a calibration ladder to convert fitness values (derived from barcode sequencing counts) to functional values (the expression level of genes regulated by the CymR transcription factor).

The normalization controls were selected by incubating a portion of the CymR library culture overnight in M9 media containing glycerol and 50 µg/mL ampicillin (M9-Gly-Amp) media (**Table S1**) with 20 ug/mL tetracycline to select for variants constitutively expressing the tetracycline resistance gene, TetA. Culture conditions were: 25 mL culture in 125 mL baffled culture flask, 37 °C with 300 rpm shaking, overnight (~16 hours). The resulting culture was streaked on LB agar plates supplemented with ampicillin and several independent colonies were picked. Two colonies were verified to have the correct plasmid sequence (using Oxford Nanopore sequencing), with CymR mutants containing insertions early in the coding sequence (one colony with I20_A21insQ, the second with E67_W68insQ). Glycerol freezer stocks for each of the two normalization controls were prepared and used in the GROQ-Seq assay as described below.

**Table S1:** Composition of M9-Gly-Amp media

| Media Component | Concentration |
|---|---|
| $KH_2PO_4$ | 3 g/L |
| $Na_2HPO_4$ | 6.78 g/L |
| NaCl | 0.5 g/L |
| $NH_4Cl$ | 1 g/L |
| $CaCl_2$ | 0.1 mmol/L |
| $MgSO_4$ | 2 mmol/L |
| Glycerol | 0.4 % |
| Casamino acids | 2 g/L |
| Ampicillin | 100 µg/mL |

The calibration ladder variants were picked as random selections from the CymR library. A portion of the library was diluted on LB agar plates supplemented with ampicillin. Twenty colonies were picked and 15 of them were verified to have the correct plasmid assembly (using Oxford Nanopore sequencing). A mixed glycerol freezer stock was prepared with equal proportions of each of those 15 calibration controls and then used with the GROQ-Seq assay (see below). Flow cytometry was used to measure the dose-response of each calibration control with the three ligands (perillic acid, perillyl alcohol, and limonene), and those cytometry results were used as the calibration datasheet to convert the GROQ-Seq results from fitness to function (see below).

GROQ-Seq DMS Assays.

The growth-based quantitative DMS assay was performed using a protocol similar to the one used for previous work with the lac repressor (LacI) and the RamR transcriptional regulator[1,2]. To reduce the number of unique barcoded plasmids in the CymR library so that long-read DNA sequencing could be used to measure the CymR coding sequence for each barcode, the transformed CymR library was bottlenecked to approximately 200,000 colony forming units following the version of the following protocol requiring “Basic Microbial Culture Equipment”: https://www.protocols.io/view/library-bottlenecking-protocols-x54v9227pl3e/v3. Several glycerol stocks were prepared from the bottlenecked library and stored at -80 C until use.

After bottlenecking, long-read (Oxford Nanopore) sequencing was used to determine the DNA barcode and CymR coding sequence for each plasmid in the library. First, one aliquot of the library freezer stock was thawed and grown to high density in 200 mL of M9-Gly-Amp media (culture volume equally split between two 250 mL baffled culture flasks, overnight incubation at 37 °C with 300 rpm shaking). Then, plasmid DNA was extracted from the overnight cultures and digested with NotI to linearize the CymR library plasmid. The extracted plasmid was then treated with calf intestinal alkaline phosphatase (CIP) and the full-length linear fragment was purified with gel extraction. Linearized DNA was sent to the sequencing provider (PlasmidSaurus) and 30 gigabases of Oxford Nanopore sequencing data were obtained. Long-read sequencing data was analyzed using the bioinformatic workflow described in Tack *et al.*[1]. The CymR coding sequence was determined for a total of 122,783 distinct barcoded plasmids, with a median coverage of 17 reads per barcode.

To prepare cultures for the GROQ-Seq measurement, one aliquot of the bottlenecked CymR library freezer stock (1 mL) was combined with 99 mL M9-Gly-Amp media and incubated in a 250 mL baffled flask at 37 C with 300 rpm shaking for approximately 17 hours. At the same time, 5 mL cultures were prepared in M9-Gly-Amp using scrapings from the glycerol stocks for the two normalization controls and the mixed calibration ladder variants. These cultures were also incubated in 15 mL snap-cap culture tubes at 37 °C with 300 rpm shaking for approximately 17 hours. After the 17 hour incubation, the following were combined in a new 250 mL baffled culture flask: 48 mL CymR library culture, 0.5 mL of each normalization culture control culture, 1 mL of the mixed calibration ladder culture, 49 mL M9-gly-amp, and 1 mL DMSO.
6 hours and 15 minutes later, the new mixed culture was loaded into an integrated automated culture and measurement system for the actual GROQ-Seq assay.

In the automated culture system, the mixed culture was sequentially diluted and incubated in a series of five 96-well growth plates (4titude cat. no. 4ti-0255, square wells, 1.1 mL per well capacity):

- Growth plate 1: 50 µL of the mixed culture diluted into 450 µL media in each well of the plate, then incubated at 37 °C for 12 hours and 5 minutes.
- Growth plate 2: 10 µL from each well of growth plate 1 diluted into the corresponding well of growth plate 2 with 490 µL media plus added ligand, then incubated at 37 °C for 3 hours and 5 minutes.
- Growth plates 3-5: 50 µL from each well of the previous plate diluted into the 450 µL media plus added ligand and tetracycline, then incubated at 37 °C for 3 hours and 5 minutes.

All media and ligand stocks used to prepare the growth plates included DMSO at a final concentration of 1%. Growth plate 2 included dilution series gradients for each of the three ligands, with six non-zero concentrations and maximum concentrations of 2500 µmol/L, 500 µmol/L, and 500 µmol/L for perillic acid (PA), perillyl alcohol (Per-OH), and S-limonene (S-Lim), respectively. Growth plates 3-5 included the same ligand dilution-series gradients, plus the addition of 5 µg/mL tetracycline (tet) to some of the wells. The plate layout for the first two rows of growth plates 3-5 is shown in **Table S2.**

**Table S2:**

| | 1 | 2 | 3 | 4 | 5 | 6 | 7 | 8 | 9 | 10 | 11 | 12 |
|---|---|---|---|---|---|---|---|---|---|---|---|---|
| **A** | media + tet | media | media + tet | 625 mol/L PA | 125 µmol/L Per-OH | 125 µmol/L S-Lim | 25.6 µmol/L PA, + tet | 64 µmol/L PA, + tet | 160 µmol/L PA + tet | 400 µmol/L PA + tet | 500 µmol/L PA + tet | 2500 µmol/L PA + tet |
| **B** | 500 µmol/L Per-OH + tet | 200 µmol/L PER-OH + tet | 80 µmol/L PER-OH + tet | 32 µmol/L PER-OH + tet | 12.8 µmol/L PER-OH + tet | 5.12 µmol/L PER-OH + tet | 5.12 µmol/L S-LIM + tet | 12.8 µmol/L S-LIM + tet | 32 µmol/L S-LIM + tet | 80 µmol/L S-LIM + tet | 200 µmol/L S-LIM + tet | 500 µmol/L S-LIM + tet |

For each growth plate, the same layout was repeated 3 additional times in rows C-H (four replicate wells for each growth condition), and for growth plate 2, the layout was the same as for growth plates 3-5 without the tetracycline. Before incubation, each growth plate was sealed with a gas permeable membrane (4titude, cat. no. 4ti-0598) to provide reproducible growth across all 96 well in each plate[3]. Each growth plate was incubated in a multi-mode plate reader (Biotek Neo2SM); the optical density (OD600) and fluorescence (569 nm excitation, 593 nm emission) were measured every 5 minutes; and continuous shaking was applied between OD and fluorescence reads (double-orbital at

807 cycles per minute). For growth plates 2-5, after 50 µL was removed from each well and transferred to the next growth, the remaining culture from the four replicate wells for each condition were combined, each into a single well in a deep well plate (Eppendorf cat. no. 951033405) to give 24 culture samples.

Plasmid DNA was then extracted from those 24 samples each culture using the automated protocol described here: https://www.protocols.io/view/automation-protocol-for-plasmid-dna-extraction-fro-8epv514yjl1b/v1.

For each growth plate, the plasmid DNA was then prepared for barcode sequencing following this protocol: https://www.protocols.io/view/automation-protocol-for-dna-barcode-sequencing-lib-81wgbp3jnvpk/v1.

The resulting barcode sequencing libraries were sent to a sequencing provider for Illumina sequencing (paired-end 150 bp). Two lanes of NovaSeq X+ data were obtained: the sample for the first lane was a balanced mixture of DNA for all four time points; the sample for the second lane was a balanced mixture of DNA for just the last three time points (growth plates 3-5). Barcode sequencing data were analyzed following workflows similar to those described for the previous work with LacI[1]. Approximately 1.3 billion barcode reads were obtained that passed all the quality checks across the 96 barcode sequencing samples (24 culture conditions × 4 time points).

The read count for each barcoded plasmid in each sample was normalized by the read count for one of the normalization controls to correct for differences in DNA extraction and PCR efficiency across different samples. The change in the normalized read count over the four time points was used to determine the fitness associated with each barcoded plasmid in each of the 24 culture conditions. The fitness values for the 15 calibration ladder plasmids were used with the flow-cytometry-based calibration datasheet (see above) to create a calibration curve for the GROQ-Seq dataset (**Figure. S3**).

The calibration curve was used to estimate the function for every barcoded CymR variant with two models of the CymR dose response: a Hill equation model and a Gaussian process model. The Hill equation model provided estimates for the parameterized dose response (basal output, $G_0$; output at the maximum concentration of each ligand, the sensitivity for each ligand, $EC_{50,lig}$; and effective cooperativity for each ligand, $n_{lig}$). The Gaussian process model provided an estimate of the dose response for each ligand as a smooth non-parametric curve. Model fitting was performed using Bayesian inference with Markov chain Monte Carlo (MCMC) using the cmdstanpy interface to Stan[4,5].

After fitting the data to determine the dose-response curves for each barcoded plasmid, the data were sorted into groups of barcoded plasmids with the same CymR amino acid sequence. For each group of barcoded plasmids (i.e., each CymR amino acid sequence), outliers (with dose-response that differed significantly from the group mean) were dropped, and the barcode read counts for the remaining plasmids were merged. 7271 distinct CymR amino acid sequences were found with a single substitutions, insertions, or deletions, with a median of 5 barcodes per amino acid sequence. For the wild-type CymR, the ten barcoded plasmids with the highest total read counts were left unmerged to provide a set of controls to assess measurement reproducibility, and data for the remaining wild-type amino acid plasmids were merged. After merging the barcode read counts for each amino acid sequence, the analysis steps from read-count normalization to dose-response fitting were repeated using the merged data to produce the results presented in the main text and in SI file 1.

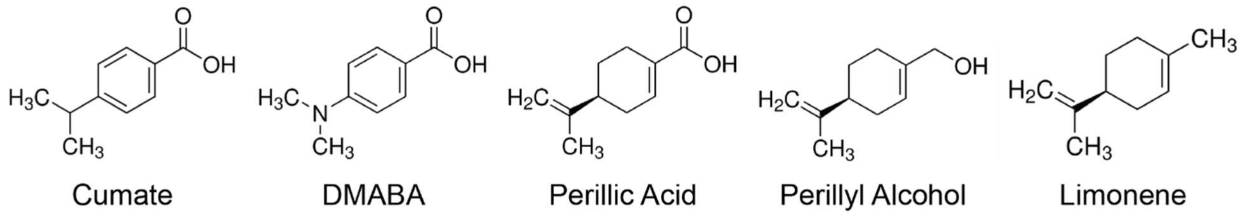


**Figure S1:** Chemical structure of five small-molecule ligands used in this work. CymR$^{AM}$ natively responds to Cumate, DMABA, and perillic acid, but is less sensitive to perillic acid. CymR$^{AM}$ does not natively respond to perillyl alcohol or limonene.

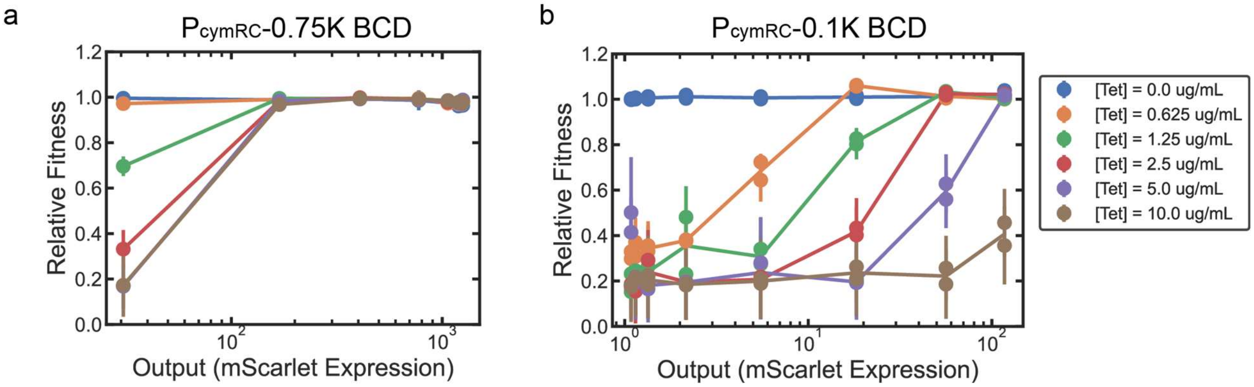


**Figure S2:** GROQ-Seq circuit tuning. The relative fitness values, plotted along the y-axis, were measured across a range of different tetracycline concentrations (indicated in the figure legend) and different induction levels (with DMABA) using the protocol described at: https://dx.doi.org/10.17504/protocols.io.8epv5xye4g1b/v1. The gene expression output, plotted along the x-axis, was measured at different induction levels (with DMABA) using flow cytometry following the protocol described at: https://dx.doi.org/10.17504/protocols.io.dm6gpzwx8lzp/v1. Results are shown for two different CymR circuits. (a) Fitness and fluorescence output values when using the 0.75K BCD translational control element to regulate translation of the TetA and RFP proteins. (b) Fitness and fluorescence output when the circuit contains an ultra-weak 0.1K BCD translational control element to regulate translation of the TetA and RFP proteins.

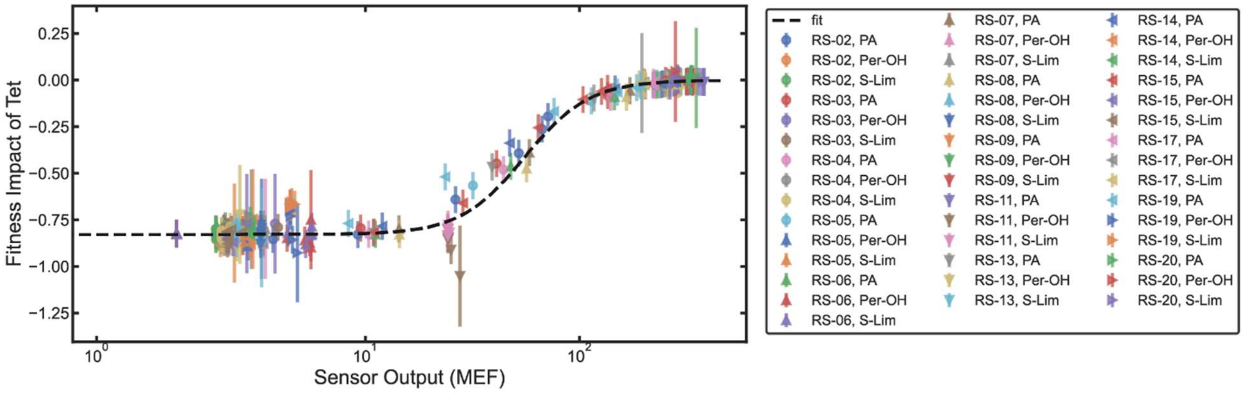


**Figure S3:** Calibration variants for GROQ-Seq. 19 calibration variants (RS-02 – RS-20) were measured using the GROQ-Seq platform for fitness against perillic acid (PA), perillyl alcohol (Per-OH) and limonene (Lim). The fitness impact of tetracycline as a function of the sensor output is plotted, with a line of best fit shown as a black dashed line.

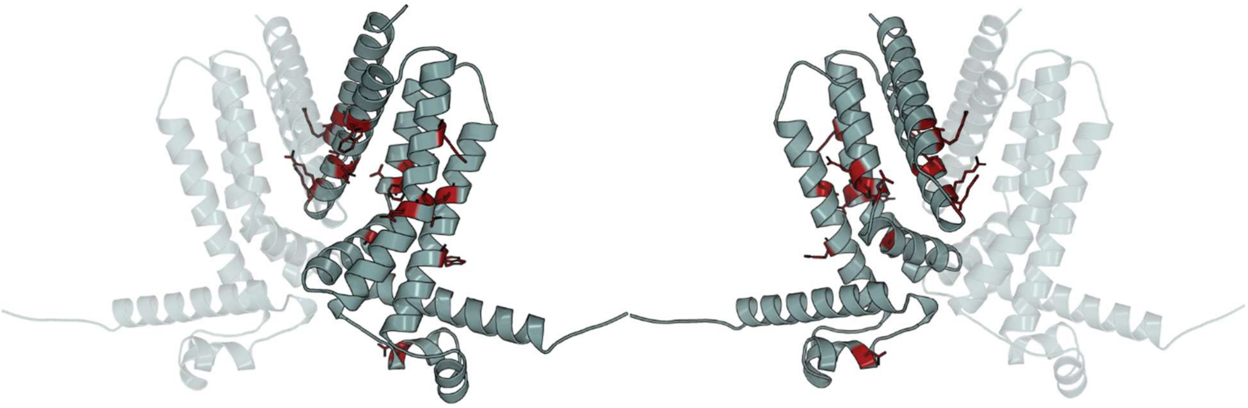

**Figure S4:** Structural representation of CymR with residues highlighted in red where more than 20% of mutations result in a phenotype where CymR has a low basal signal when ligand is absent (G0), indicative of DNA repression, as well as a low fold change between no ligand and 2500 µmol/L PA (G2500/G0). Side chains for the selected residues are shown.

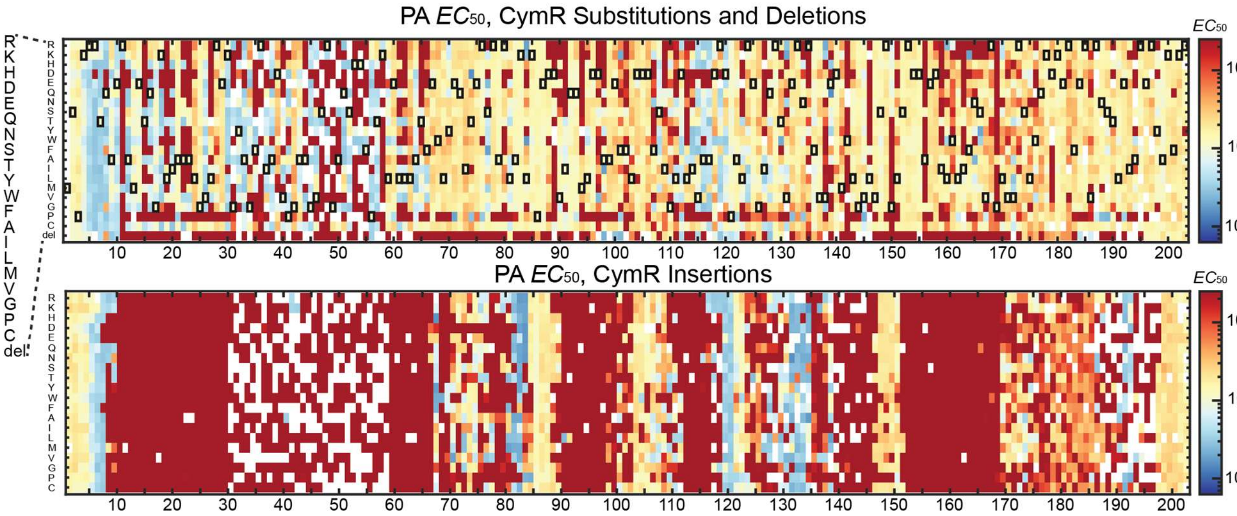


**Figure S5**: Heatmaps showing the $EC_{50}$ with perillic acid from the GROQ-Seq measurement for all tested variants. Dark red indicates variants which showed no response to PA across all tested concentrations. Wildtype residues are shown in black boxes. White spaces indicate variants which did not appear in the library.

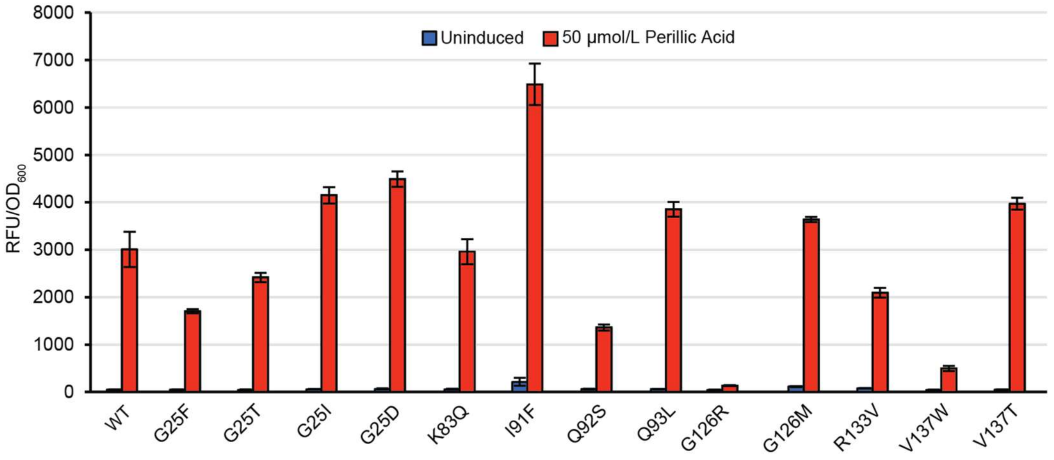


**Figure S6:** Normalized fluorescence of CymR variants that were identified via GROQ-Seq to have improved fold change in response to perillic acid with and without 50 µmol/L perillic acid as compared to the WT CymR$^{AM}$.

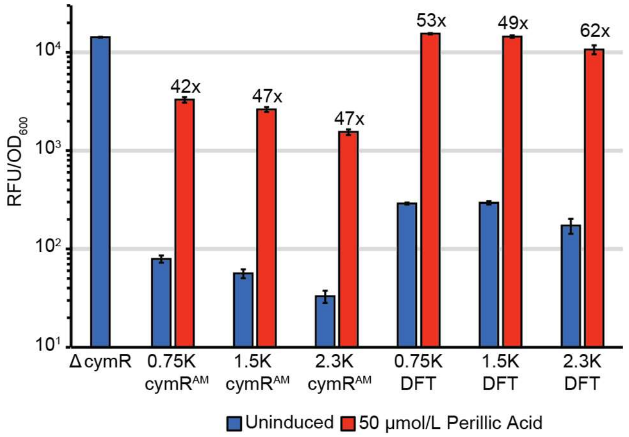


**Figure S7:** Normalized fluorescence for the WT CymR$^{AM}$ and DFT triple mutant expressed from BCD translational control elements with increasing strength and both uninduced and induced with 50 µmol/L perillic acid.

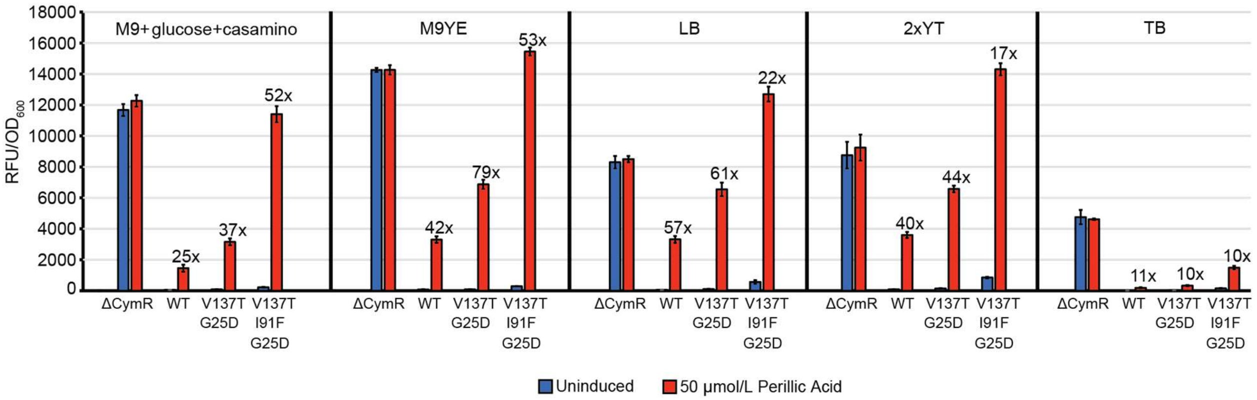


**Figure S8:** Normalized fluorescence for three variants of CymR$^{AM}$ in five different media conditions. The WT CymR$^{AM}$ protein, a double mutant with V137T and G25D mutations, and a triple mutant containing an additional mutation at I91F were tested in M9YE, LB, M9+glucose+casamino acids, 2xYT, and TB media, both uninduced and with 50 µmol/L perillic acid. Fold changes between uninduced and induced with 50 µmol/L perillic acid values are shown above the data bars.

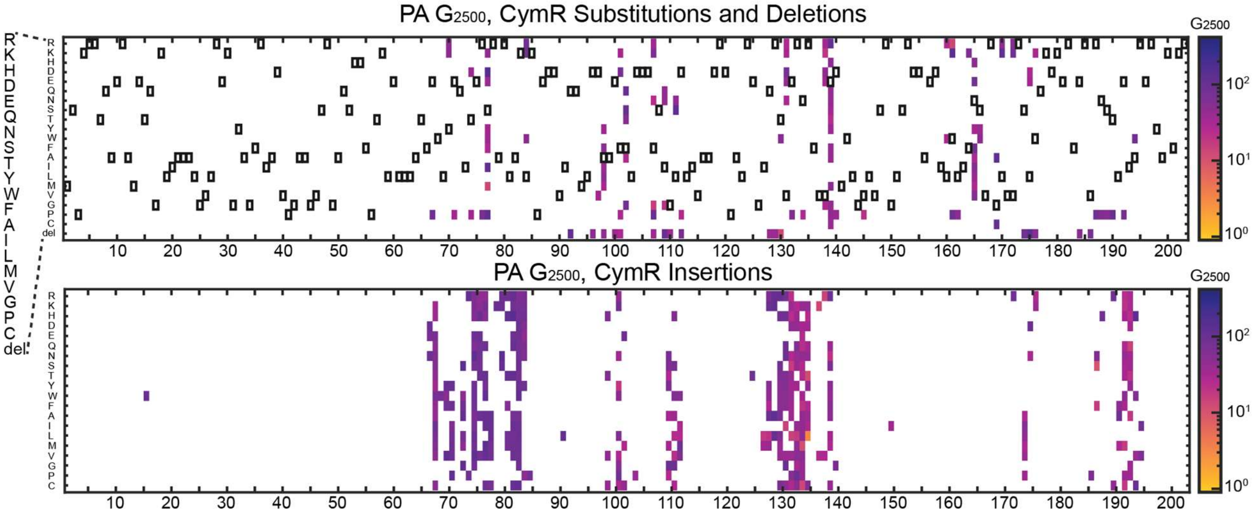


**Figure S9:** The transcription signal at 2500 μmol/L perillic acid induction, gated to show only CymR variants with a 97.5% confidence of having an inverted phenotype. The top heatmap shows substitutions and deletions, and has black boxes indicating the wildtype amino acid at each position. The bottom heatmap shows CymR variants with an insertion mutation, where there are no wildtype variants.

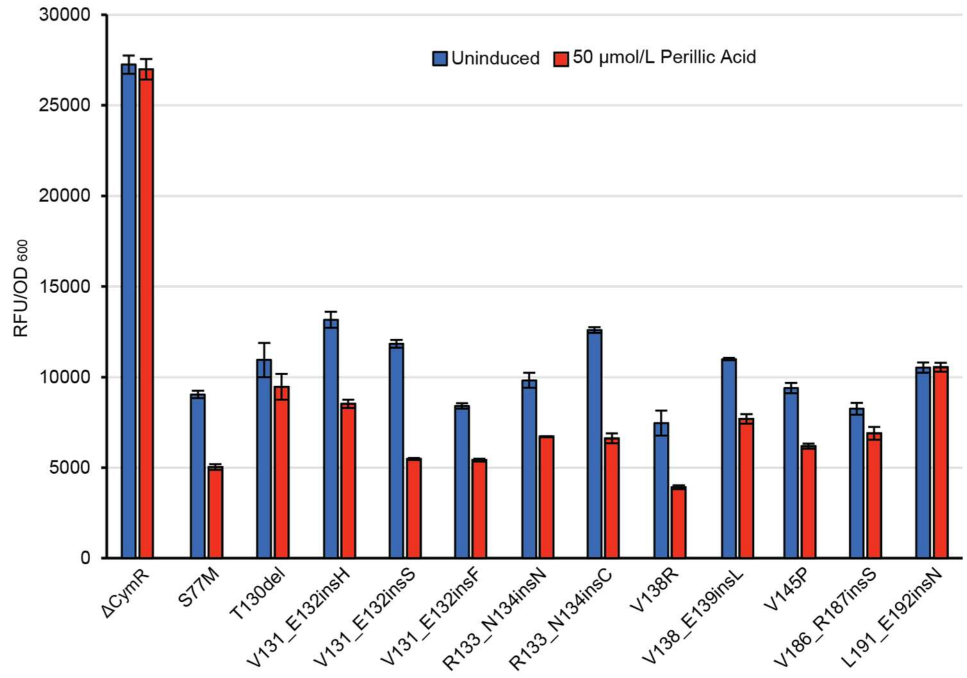


**Figure S10:** Normalized fluorescence for 12 different variants of $CymR^{AM}$ induced with 0 or 50 µmol/L perillic acid. Variants were selected following GROQ-Seq for further investigation of the inverted phenotype.

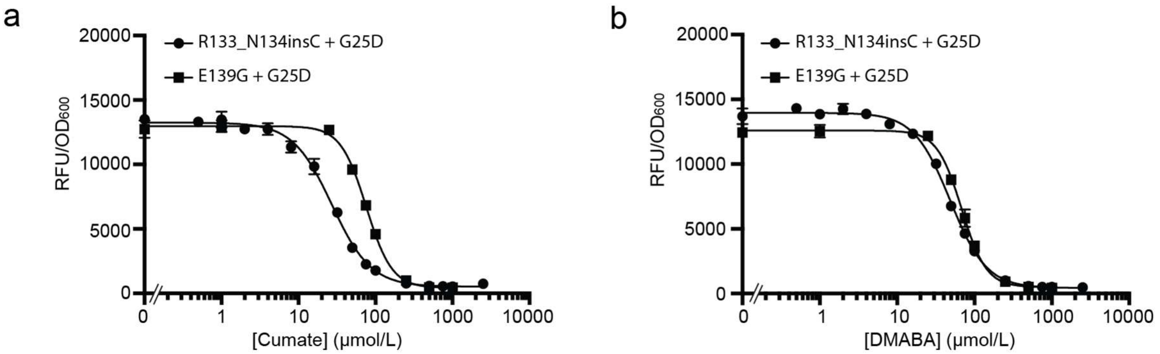


**Figure S11:** Induction curves for the best inverted CymR$^{AM}$ variant discovered in this work compared to the E139G +G25D variant. (a) CymR$^{AM}$ variants R133N134insC + G25D and E139G + G25D variant induced with cumate over a range of concentrations. Line represents the logarithmic curve of best fit. (b) CymR$^{AM}$ variants R133N134insC + G25D and E139G + G25D variant induced with DMABA over a range of concentrations. Line represents the logarithmic curve of best fit.

**Table S3:**

| | |
|---|---|
| RS_name | The name of the calibration variant (if applicable) |
| mutation_positions | A list of positions that are mutated |
| cymR_amino_mutations | Number of amino acid mutations |
| has_amino_indel | Boolean indicating whether or not the CymR amino acid sequence has one or more indels |
| mutation_codes | List of mutations |
| cymR_amino_seq | The amino acid sequence of the CymR variant |
| log_g0 | Log10 of the basal output from the CymR sensor, from the Hill model fit |
| log_g0_err | Uncertainty of the log_g0 |
| GP_log_g0 | Log10 of the basal output from the CymR sensor, from the Gaussian Process model fit |
| GP_log_g0_err | Uncertainty of the GP_log_g0 |
| log_ginf_{ligand} | Log10 of the saturated output when induced with the ligand from the CymR sensor |
| log_ginf_{ligand}_err | Uncertainty of the log_ginf_{ligand} |
| log_ec50_{ligand} | Log10 of the EC50 of the dose-response with {ligand} |
| log_ec50_{ligand}_err | Uncertainty of the log_ec50_{ligand} |
| sensor_n_{ligand} | The effective cooperativity of the dose-response (Hill coefficient), with {ligand} |
| sensor_n_{ligand}_err | Uncertainty of the sensor_n_{ligand} |
| log_ginf_g0_ratio_{ligand} | Log10 of the ratio of the saturated to basal output, with {ligand} |
| GP_log_g{conc}_{ligand} | Log10 of the output at the maximal induction with {ligand}, where {conc} is the max concentration measured, from the Gaussian Process model fit |
| GP_log_g{conc}_{ligand}_err | Uncertainty of the GP_log_g{conc}_{ligand} |
| GP_log_g{conc}_g0_ratio_{ligand} | Log10 of the ratio of the maximally induced and basal output from the CymR sensor, where {conc} is the max concentration |

| | |
|---|---|
| | measured, from the Gaussian Process model fit |
| GP_log_g{conc}_g0_ratio_{ligand}_err | Uncertainty of the GP_log_g{conc}_g0_ratio_{ligand} |
| total_counts | The total number of barcode reads used in the data analysis for the CymR variant |
| total_counts_plate_2 | The total number of barcode reads for growth plate 2 (the 1st time point |
| nanopore_count | The total number of Nanopore reads for the CymR variant |
| amino_type_count_0 | The number of distinct barcodes for the CymR variant, before dropping outliers |
| amino_type_count_f | The number of distinct barcodes for the CymR variant, after dropping outliers (i.e., the number of barcodes merged for data analysis) |

**Table S4:** A list of all positions that were indicated to have an induction curve with both a negative slope and a positive slope at different points.

| |
|---|
| Y70E71insN<br>Y70E71insC |
| E71Q72insG |
| I73T74insN<br>I73T74insT |
| E75R76insK |
| R80L81insM |
| L81A82insQ |
| A82P |
| A82K83insE<br>A82K83insF<br>A82K83insH<br>A82K83insN<br>A82K83insQ<br>A82K83insT |
| K83L84insD<br>K83L84insG<br>K83L84insH<br>K83L84insN<br>K83L84insQ<br>K83L84insS<br>K83L84insT |
| L84W |
| F102L103insY |
| D112del |
| I114G |
| E132R133insQ |
| V138E139insY |
| E139H |
| D158R |
| R185V186insE |
| V186R187insH<br>V186R187insR |
| S189T190insF |
